# nERdy: network analysis of endoplasmic reticulum dynamics

**DOI:** 10.1101/2024.02.20.581259

**Authors:** Ashwin Samudre, Guang Gao, Ben Cardoen, Bharat Joshi, Ivan Robert Nabi, Ghassan Hamarneh

## Abstract

The endoplasmic reticulum (ER) comprises smooth tubules, ribosome-studded sheets, and peripheral sheets that can present as tubular matrices. ER shaping proteins determine ER morphology, however, understanding their role in tubular matrix formation requires reconstructing the dynamic, convoluted ER network. Existing reconstruction methods are sensitive to parameters or require extensive annotation and training for deep learning. We introduce nERdy, an image processing based approach, and nERdy+, a D4-equivariant neural network, for accurate extraction and representation of ER networks and junction dynamics, outperforming previous methods. Comparison of stable and dynamic representations of the extracted ER structure reports on tripartite junction movement and distinguishes tubular matrices from peripheral ER networks. Analysis of live cell confocal and STED time series data shows that Atlastin and Reticulon 4 promote dynamic tubular matrix formation and enhance junction dynamics, identifying novel roles for these ER shaping proteins in regulating ER structure and dynamics.

## 1 Main

The endoplasmic reticulum (ER) is the largest membrane-bound organelle, spanning the cytoplasm from the nucleus to the plasma membrane as a continuous membrane network. The ER serves as a major site of protein synthesis, folding, and sorting, as well as various cellular functions including stress response, Ca2+ storage, and lipid metabolism [1]. The peripheral ER forms an extended tubular network connected to more densely labeled sheet-like regions; relative expression of the sheet-inducing lumenal spacer protein CLIMP-63 and membrane curvature-inducing Reticulon 4 determines the abundance of peripheral ER tubules and sheets, as well as thickness of the tubules and sheets [2–5]. The Atlastin (ATL) GTPase promotes the homotypic fusion of ER tubules and the formation of tripartite junctions in peripheral ER tubular networks [6, 7]. LunaPark localizes to tripartite ER junctions and interacts with Atlastin and Reticulon 4 to regulate the extent of the tubule and junction formation [8, 9]. Atlastin regulation of ER junction dynamics, but not LunaPark, has been shown to impact microtubule organization [10]. While LunaPark stabilizes junctions on the order of minutes [11], ER network dynamics can be far more rapid [12–15].

High-speed studies using STED and single molecule super-resolution microscopy showed that Reticulon 4 and CLIMP-63 control the size and dynamics of lumenal nanodomains along peripheral ER tubules [13, 14, 16]. Identification of CLIMP-63 as a sheet-inducing ER shaping protein was based in large part on the identification of sheet-like structures in the cell periphery by diffraction-limited confocal microscopy [3]. However, subsequent analysis by grazing incidence SIM combined with light sheet imaging, increasing both the spatial and temporal resolution, showed that peripheral sheets are actually composed of a dense matrix of tubules [15]. The tubular matrix corresponds to the convoluted tubular networks observed by electron microscopy (EM) for smooth ER and known as the site of viral replication [17–19]. Deep learning analysis of 3D super-resolution microscopy identified the tubular matrix as a distinguishing feature of ER in Zika infected cells [20]. However, understanding dynamic ER behavior in peripheral tubular networks versus denser tubular matrices requires measures to accurately reconstruct ER networks, define the distinct peripheral ER regions, and capture junction dynamics.

Several methods have been developed to analyze the ER structure. The morphology of plant ER structure has been recreated using a network and graph-theoretical approach, however, these approaches do not address the dynamic rearrangement of ER networks [21–25]. A study of lumenal particle dynamics suggests that exploration time can be a measure of ER network connectivity [26]. ER dynamics and curvature properties were analyzed using point tracking, contour tracking, and Fourier decomposition methods [27]. Some other methods such as phase-congruency analysis were used for the segmentation of tubules and sheets (cisternae) [28]. In [29], the authors developed a deep residual-network model, ER-net, for the segmentation of ER. In follow-up work, ERnet-v2 [30], the authors used a Swin Transformer-based model to extract tubules, sheets, and sheet-based-tubules (SBTs) from the ER structure and provided quantitative measures to understand the topology of the ER network in a supervised manner. Garcia et al. [31] developed a pipeline to quantitatively measure the areas of rough and smooth ER to assess the impact of different pharmacological perturbations on ER morphology. Analysis of ER dynamics is non-trivial because of the low signal-to-noise ratio and highly variable fluorescence intensity distribution over space and time [32].

Here, we provide two approaches for the reconstruction and analysis of ER structure. Our first approach, nERdy, employs classical image and graph processing algorithms for geometrical structure analysis [33], offering a computationally efficient solution with minimal parameter tuning. Our second approach, nERdy+, is an equivariant encoder-decoder neural network that enhances the accuracy of ER structure extraction, eliminating the need for parameter tuning in morphological image processing operations. The equivariant architecture naturally captures data symmetries and transformations, improving robustness against perturbations and enhancing generalization. Notably, our analysis of junction dynamics reveals that the ER shaping proteins Reticulon 4 and Atlastin, boost the dynamics of Isolated peripheral ER junctions and facilitate the formation of tubular matrices. Comparative evaluations against ERnet and ERnet-v2, conducted on manually annotated ground truth image sequences, demonstrate that both nERdy and nERdy+ significantly outperform ERnet-v2 in segmentation metrics as well as graph metrics.

## 2 Results

### 2.1 Dataset details

To study the impact of ER shaping proteins on peripheral ER networks, we conducted high-speed confocal time-lapse imaging (25 Hz, 100 frames) of peripheral regions of interest (ROIs) in COS-7 and HeLa cells. Both COS-7 and HeLa cells were transfected with the lumenal ER reporter, ERmoxGFP, as well as Reticulon 4, CLIMP-63, and mCherry-tagged Atlastin. The COS-7 dataset encompasses a total of 117 timelapse series, obtained from three biological replicates. It includes 31 Control sequences featuring ERmoxGFP, 26 sequences with dual-channel ERmoxGFP/Atlastin-mCherry, and 31 sequences each for ERmoxGFP/CLIMP-63 and ERmoxGFP/Reticulon 4. Representative images of the time-lapse series acquired are shown in Figure 1. The HeLa dataset consists of a total of 119 time-lapse series with 39 Control sequences, 27 Atlastin sequences, 25 CLIMP-63 sequences and 28 Reticulon 4 sequences. While the ER is inherently a continuous network, time-lapse imaging of the ERmoxGFP labeled ER showed frequent discontinuities in the network and irregularities across successive frames. Time frames 25, 30, and 35 of ERmoxGFP label in a sequence from an Atlastin transfected cell show the formation and breakage of tubules over time (Extended Data Figure 1). We also utilize the STED time-lapse series (25 Hz, 100 frames) of COS-7 cells transfected with ERmoxGFP alone or ERmoxGFP and either Reticulon 4-mCherry or CLIMP-63-mCherry [13].

**Fig. 1.**
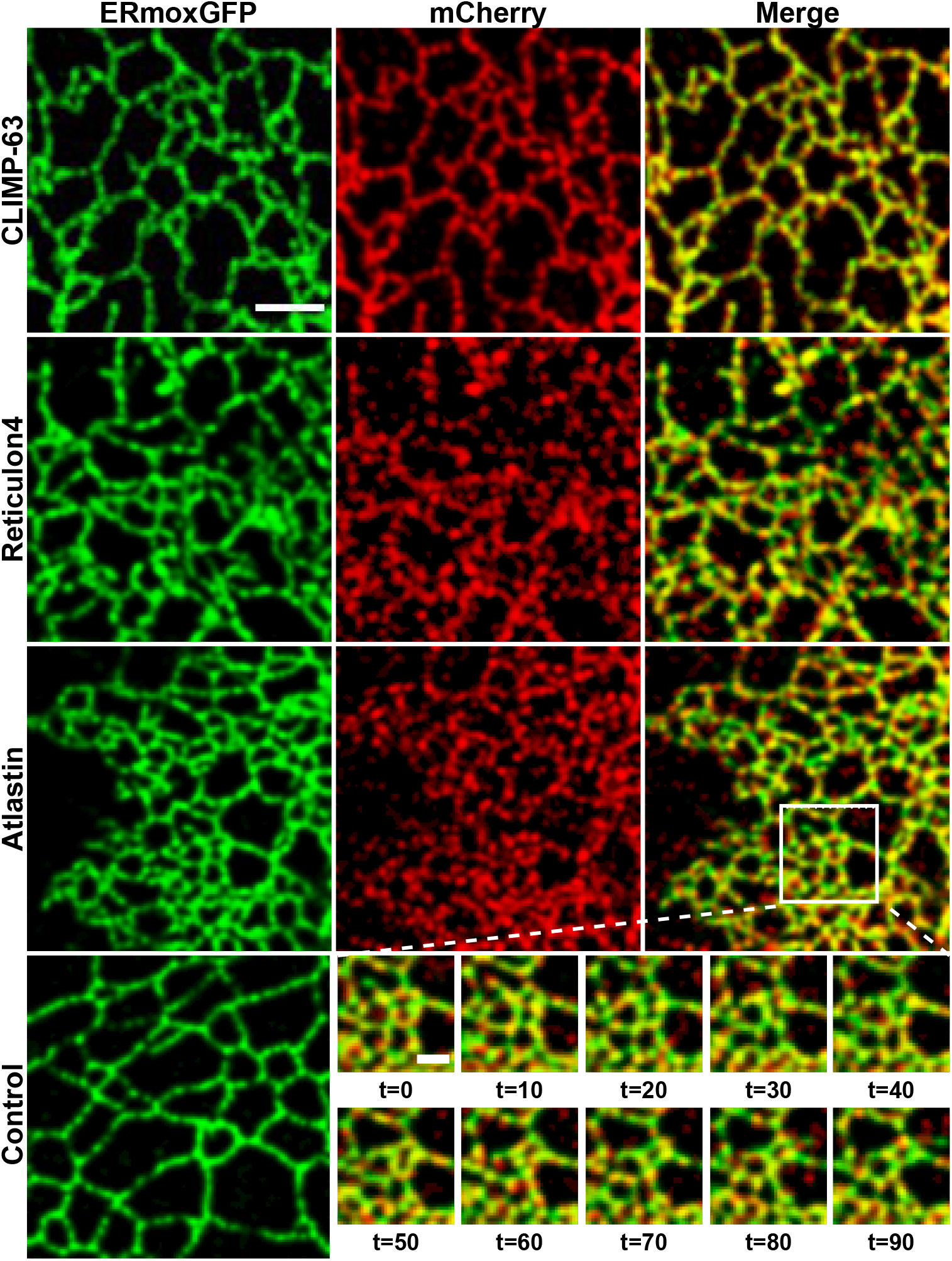
Peripheral ER time-lapse confocal imaging. COS-7 cells were transfected with ERmoxGFP (green) along with either CLIMP-63 (row 1), Reticulon 4 (row 2), or Atlastin (row 3) tagged with mCherry (red; column 2), and peripheral regions of interest (ROI) imaged by time-lapse microscopy (25 Hz, 100 frames in total). The last column denotes the merge of both the ERmoxGFP and mCherry channels. The bottom row displays the ERmoxGFP image of Control cells transfected with ERmoxGFP as well as insets (boxed region) of an Atlastin-mCherry co-transfected cell at 10-frame intervals. Scale bar: 3 *µ*m

### 2.2 Computational Analysis

To analyze the dynamic ER network and junctions in these time-lapse series, we developed a computational pipeline encompassing: 1) Segmentation of tubular ER structure; 2) Extraction of junction dynamics from temporal frames via graph processing; 3) Classification of low movement (‘Isolated’) and high movement (‘Overlapping’) junction regions within the ER structure.

#### 2.2.1 Segmentation of tubular ER structure

##### nERdy: Image processing pipeline for network extraction

In nERdy (Figure 2A), we aim to extract the tubular ER structure from an input frame of the time-lapse sequence. Our approach involved a series of operations to remove noise and fine structures while preserving tubular integrity. These include intensity normalization, histogram equalization, and morphological operations such as area opening, erosion, and local thresholding (see Section 4 for details). We then applied the Jerman Enhancement Filter with specific parameters [34] to amplify the tubular structure in the input. These steps yield a segmented ER structure suitable for skeletonization and network analysis. Our parameter configuration in morphological operations shows robustness across all samples.

**Fig. 2.**
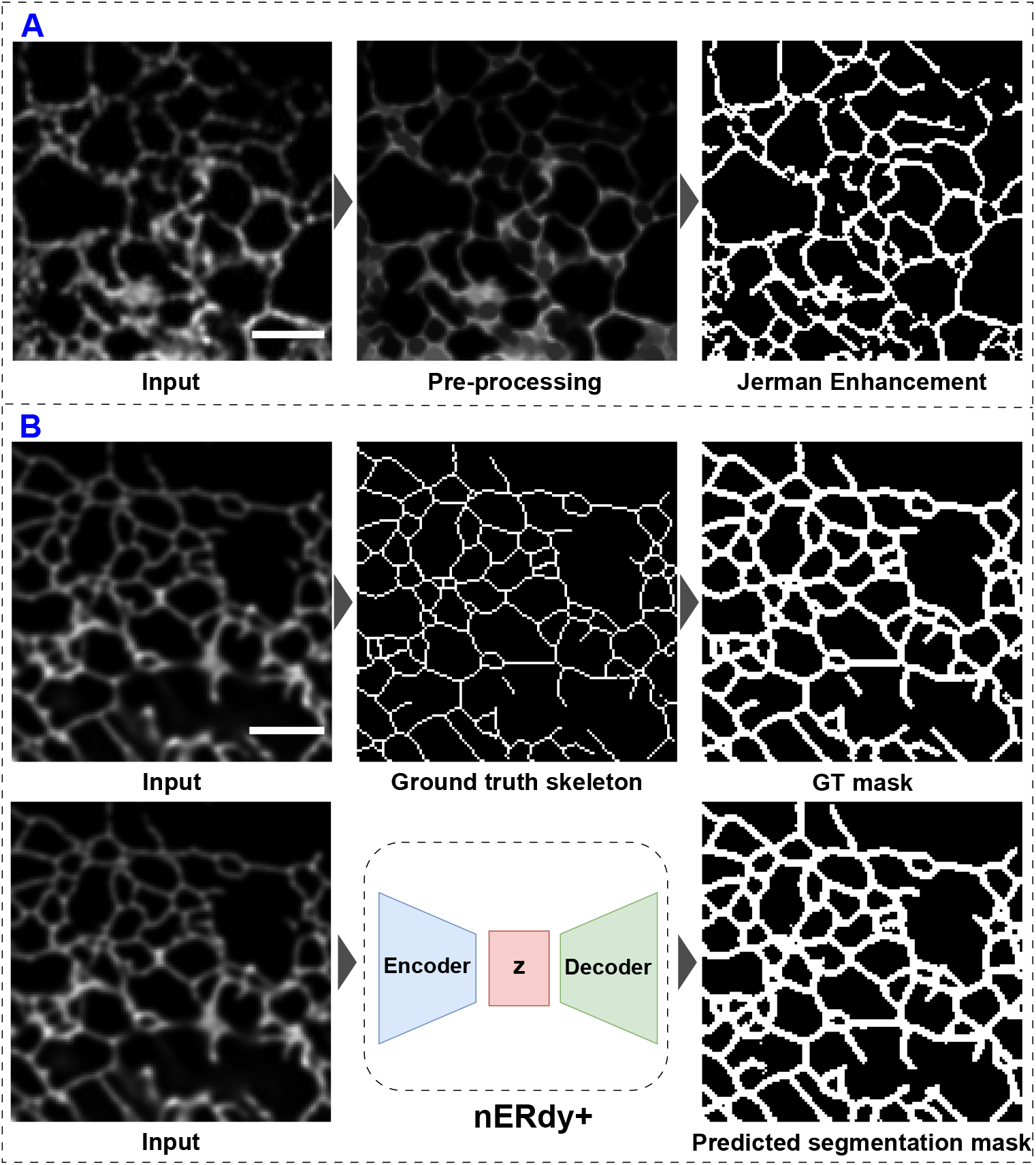
Pipelines for nERdy and nERdy+. Panel A: The input ER frame undergoes a set of morphological operations including intensity normalization, histogram equalization, area opening, erosion, and local thresholding, resulting in the preprocessed output displayed in ‘Pre-processing’. This preprocessed sample undergoes the Jerman Enhancement filtering to obtain the tubular output. Panel B: For the example input ER frame we acquire the manually annotated ground truth skeleton. To train nERdy+, a mask is created using a morphological dilation operation applied on the ground truth skeleton (‘GT mask’). After training, nERdy+ predicts a probability map for the given input. Post-processing with the Rolling Ball algorithm and Otsu thresholding generates the final segmentation output shown as ‘Predicted segmentation mask’. Scale bar: 3 *µ*m

##### nERdy+: D4-equivariant encoder-decoder network for ER tubule segmentation

To circumvent the manual parameter tuning and derive an ER structure representation learned from the data, we developed nERdy+, a deep learning based approach. nERdy+ is an encoder-decoder neural network [35, 36] (Figure 2B) trained to produce segmentation probability maps for the input ER images. To learn robust data representation from a small-scale labeled dataset (117 samples), we designed an equivariant architecture in nERdy+. Equivariance ensures that a neural network’s response should change predictably according to the input transformations. Given the significant variability present in ER images, equivariance plays a crucial role in learning the ER data representation.

We develop nERdy+ as an architecture equivariant to the Dihedral group (D4), denoting mod *π/*2 rotations and reflections (mirrored around x=0), thus providing eight views per input (Extended Data Figure 2). These symmetries guide the network towards consistent responses, ensuring consistency despite transformations. The encoder extracts hierarchical and rotationally invariant features, while the decoder reconstructs outputs in accordance with D4-equivariant representations. The output is a probability map, which undergoes the rolling ball algorithm [37] to remove low-intensity background regions. Otsu thresholding is then applied to obtain a binary segmentation. nERdy+ outperforms state-of-the-art ER segmentation approaches [29, 30], delivering faithful and accurate segmentations of the input ER structure.

#### 2.2.2 Quantitative and Qualitative evaluation of nERdy+

In both confocal and STED data, manual annotation of 100 frames per time-lapse series is not feasible. Thus, we obtain a stable representation of the ER structure sequence via the projection of all input frames onto a single frame and calculate the mean intensity values, termed as ‘Mean projection’. To train and evaluate nERdy+, we manually annotated the network structure (skeleton) of the mean projection frames across both confocal and STED time-lapse series. These ground truth skeletons are processed with morphological dilation operation to obtain segmentation masks termed as ‘GT mask’. (Figure 2B, see details in section 4.4). To train nERdy+, we utilized confocal data, employing an 80-20 split for training and validation across the 117 annotated samples along with 5-fold cross-validation. The trained model is later evaluated on completely unseen 35 annotated STED samples.

We compare the performance of nERdy and nERdy+ with three competing approaches for ER segmentation: ERnet [29], ERnet-v2 [30], and AnalyzER [28]. We obtain the segmentation maps for ERnet and ERnet-v2 using the publicly available trained models. For AnalyzER, we obtain the segmentation maps using the default set of parameters in the GUI while setting the ‘cisternae’ parameter off. This helps to primarily extract the tubular structure from the input ER samples.

We perform method evaluation at two levels: segmentation and graph measures. In segmentation, we compare the ‘GT mask’ with the predicted segmentation mask (Figure 2B) of the ER structure. For graph measures, we convert both manually annotated ground truth skeleton and extracted skeleton to a graph (see section 4.8) for comparison. Our segmentation metrics include Intersection over Union (IoU), which is also known as the Jaccard Index, Dice score, and F1-score. For two sets A and B representing ground truth mask pixels and prediction mask pixels respectively, the Jaccard Index is given as:

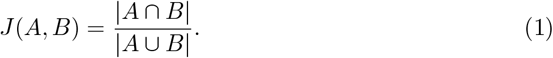

The Dice score is given as:

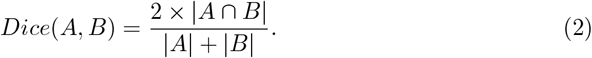

and the F1 score is given as:

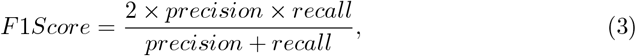

where precision is the ratio of correctly predicted segmentation mask pixels and all predicted mask pixels, whereas recall is the ratio of correctly predicted segmentation mask pixels and all the ground truth mask (‘GT mask’) pixels.

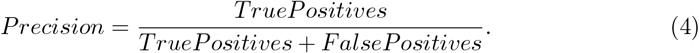

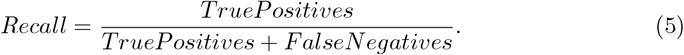

Figure 3A shows the qualitative performance of different methods. nERdy and nERdy+ both outperform the other three competing methods across the three metrics, with nERdy+ providing the highest scores (Figure 3B).

**Fig. 3.**
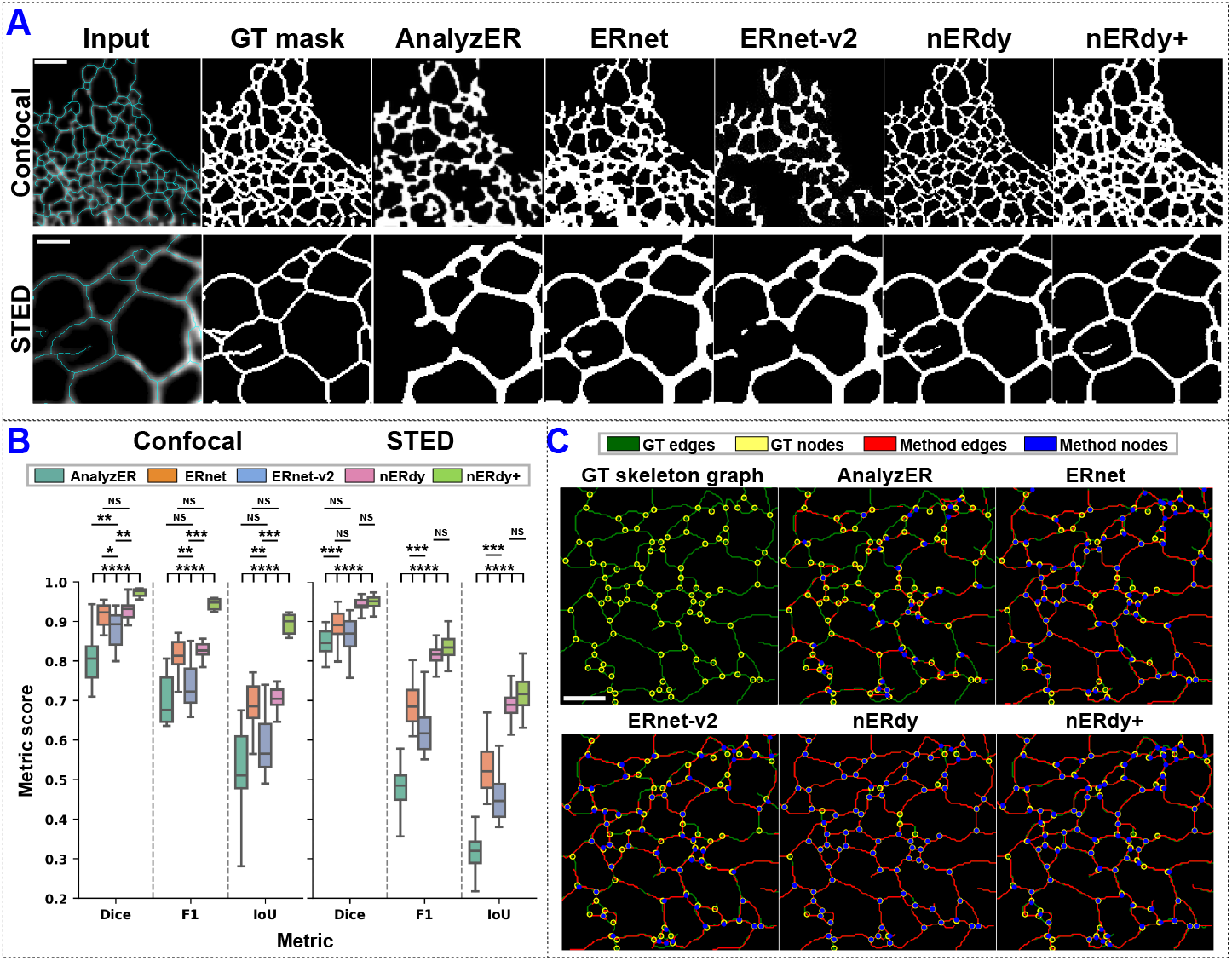
Qualitative and quantitative evaluation of different segmentation methods. A) Comparison between the different methods based on the segmentation results on one sample from confocal (row 1) and STED (row 2) time-lapse series. The cyan overlay on the input ER represents the ground truth skeleton for both confocal and STED data of COS-7 cells. ‘GT mask’ illustrates the baseline segmentation for the input samples obtained using morphological dilation of manually annotated skeletons. While ERnet and ERnet-v2 show a wider segmentation compared to the baseline, nERdy and nERdy+ show precise and superior segmentation for the input samples. Scale bar: 3 *µ*m (Confocal), 0.75 *µ*m (STED) B) Quantitative evaluation for both confocal data (*N* =21) and STED data (*N* =35). nERdy+ outperforms other methods across all the metrics in confocal data, whereas in STED data, nERdy and nERdy+ show comparable performance. Statistical significance is calculated using a two-sided Mann-Whitney U test with Bonferroni correction. P-value annotations are as follows, NS: p *<* 1.00e-02, *: 1.00e-02 *<* p ≤ 5.00e-02, **: 1.00e-03 *<* p ≤ 1.00e-02, ***: 1.00e-04 *<* p ≤1.00e-03, ****: p ≤ 1.00e-04. C) Qualitative evaluation of skeleton reconstruction. The performance is shown using an example from a confocal time-lapse series across different methods. The ground truth skeleton graph (GT skeleton graph) displays edges in green and nodes in yellow. In other cases, red edges represent the edges from the output of the selected method, and blue spots depict the nodes from the selected method. The output edges and nodes are overlaid on the ground truth graph for comparison. nERdy shows the highest precision for the location of nodes and edges but misses a few edges. nERdy+ shows less precision for edge and node locations, but captures a majority of the nodes and edges, leading to a more complete graph reconstruction. ERnet and ERnet-v2 capture a majority of nodes and edges, but still lack precision compared to nERdy and nERdy+. Scale bar: 3 *µ*m

In the case of confocal data (Figure 3B), nERdy+ outperforms previous methods across all three metrics with a Dice score of 0.96 +/-0.003, F1-score of 0.94 +/-0.005 and Jaccard index of 0.88 +/-0.01. nERdy provides the second best performance in all the metrics, followed closely by ERnet. When testing the performance on STED data (Figure 3B), we observe that nERdy+ provides the best metric values, followed closely by nERdy. The superior performance of nERdy+ across both confocal and STED data indicates its robustness and effectiveness in segmenting ER structure. The narrow confidence intervals (e.g., +/-0.003 for Dice score) suggest consistent performance of nERdy+ across different data samples, highlighting its reliability.

In Lu et al. [30], the authors present a comprehensive set of metrics to evaluate the quality of reconstructed graphs derived from ER images. These metrics include the total number of edges, the total number of nodes, assortativity coefficient, clustering coefficient, number of components, the ratio of nodes, and the ratio of edges. The assortativity coefficient measures the tendency of nodes to connect to nodes with similar degrees, while the clustering coefficient quantifies how nodes tend to cluster together, providing insights into small-scale structures. Additionally, the number of components indicates graph connectivity, and the ratios of edges and nodes quantify subgraph properties. The ratio of edges is defined as the number of edges in the largest subgraph divided by the total number of edges in the graph. Similarly, the ratio of nodes is defined as the number of nodes in the largest subgraph divided by the total number of nodes in the graph.

Here, we introduce two additional metrics for performance analysis: local efficiency and density. Local efficiency measures the immediate exchange of information within node neighborhoods, indicating well-connected neighborhoods. Density represents the ratio of actual edges to possible edges in the graph, indicating network connectivity. In our analysis, we calculate relative error compared to ground truth skeleton graphs. Considering k samples in a set, the relative error is given as:

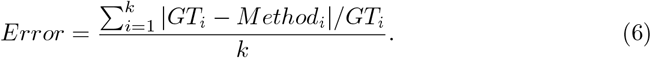

We find that nERdy+ exhibits the lowest error across the majority of metrics. For confocal data (Extended Data Figure 3), nERdy+ outperforms other methods on 7 out of 9 metrics. However, nERdy shows slightly better performance in the number of edges metric, while ERnet shows the least error in the assortativity coefficient metric. ER network can be considered as a collection of edges joined at the nodes, and thus accurate reconstruction of both the edges and nodes is essential to understand the underlying ER structure. In the case of STED data (Extended Data Figure 4), nERdy+ achieves the lowest error in 5 out of 9 metrics, while nERdy excels in three metrics: number of edges, assortativity coefficient, and clustering coefficient, showcasing its robust network reconstruction capability. ERnet performs best in the number of components metric.

In Figure 3C, we present output skeleton graphs for a CLIMP-63 sample using different methods. Ground truth skeleton graph edges are depicted in green, and junctions with degree greater than 2 are in yellow. Junctions obtained via each method are shown in blue. AnalyzER and, to a lesser extent, ERnet-v2, display incorrect reconstruction in several ER regions, illustrated by the high visibility of ground truth edges in the overlay. In contrast, nERdy, nERdy+, and ERnet produce faithful network reconstructions, with minor issues observed across different regions.

With segmentation, we assess the reconstruction capabilities of the methods, focusing on their ability to accurately identify and delineate structures within the images. On the other hand, graph measures provide additional insights into the structural characteristics of the reconstructed networks, such as connectivity patterns, degree distributions, and small-scale structures, offering a more comprehensive understanding of the underlying ER network topology.

We note that ERnet-v2 consistently underperforms compared to nERdy+, nERdy, and ERnet, mainly due to our data predominantly comprising tubules with minimal sheet presence. In contrast, ERnet-v2 specializes in segmenting tubules, sheets, and sheet-based tubules (SBTs).

#### 2.2.3 Extraction of Junction Dynamics

We use the binary segmentation output from nERdy+ and perform skeletonization [38] to represent the input object as one-pixel wide centerlines while preserving connectivity. The skeletonized structure is then converted into a graph representation, where nodes are identified based on 3-way bifurcation locations, corresponding to junctions in the tubular structure (Figure 4A). The movement of the ER structure over time leads to the emergence of dynamic junction locations in subsequent frames. As shown in Extended Data Figure 5A, an example junction appears at four adjacent but different locations in the first four frames, leading to the formation of a connected component (CC) after four time steps. To visualize the overall movement of junctions, all junction locations are plotted onto a single image, as illustrated in Figure 4B (‘Junction Projection’). Using the connected components algorithm on the binary projection frame, we identify different CCs within the input. This algorithm evaluates the connectivity of elements in the input, assigning labels to pixels based on their connectivity to neighboring pixels. The labeling process starts with 1 for the first component and increments for subsequent components, while background pixels are labeled as zero. The distinct colors representing CCs, in the ‘connected components’ panel of the last column in Figure 4B, visually illustrate this labeling process.

**Fig. 4.**
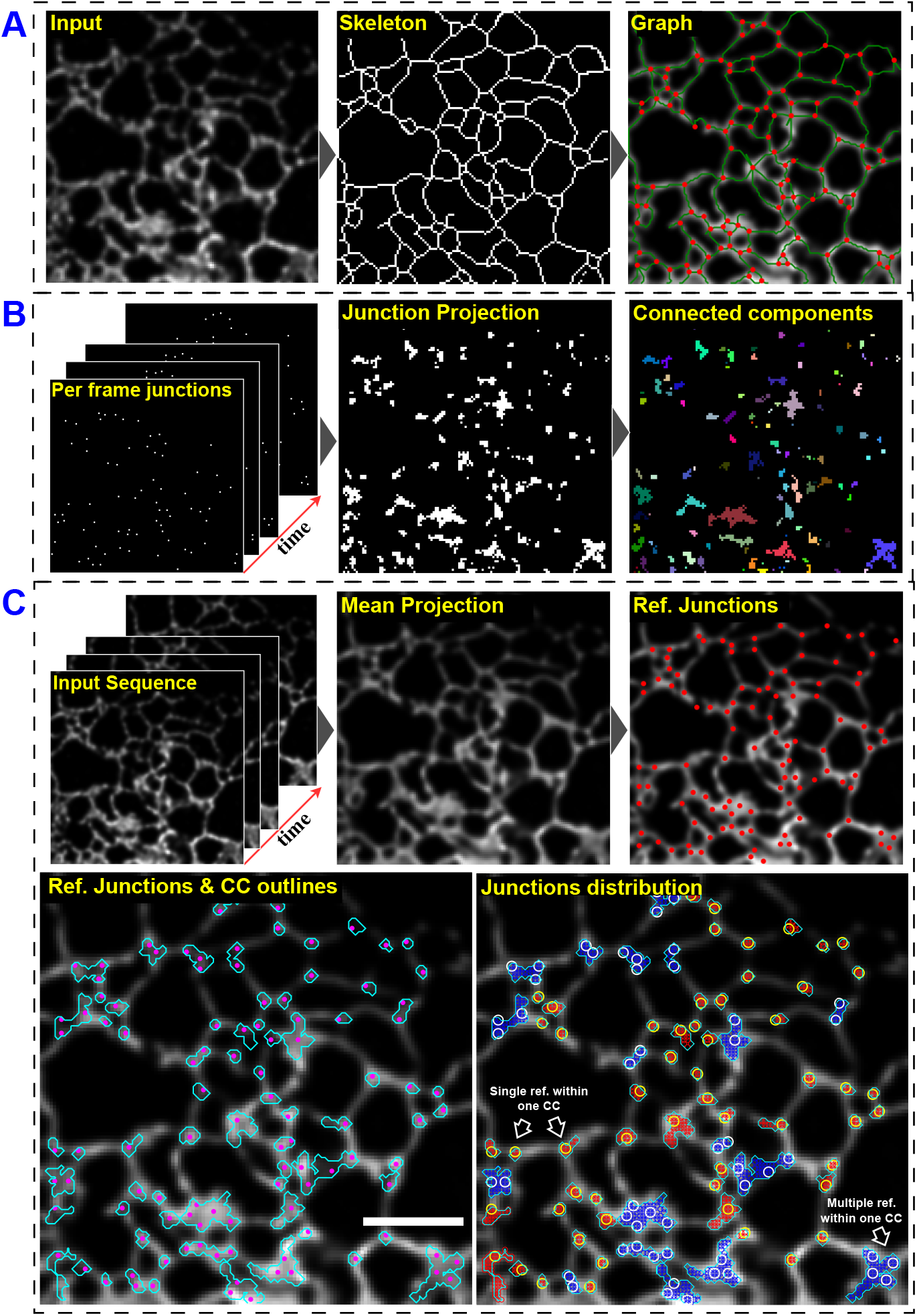
Junction dynamics extraction and classification. A) The input frame is processed with nERdy/ nERdy+ followed by skeletonization to get a skeleton. The skeleton is converted to a graph with edges depicting the tubules of the ER and nodes depicting the junctions. B) Junctions obtained for each frame in the sequence (‘Per frame junctions’) are projected onto a single frame ‘Junction Projection’. As the last step, connected component labeling is performed for the junction projection to define junction movement regions (CC) for the time-lapse series. C) Junction classification: Mean projection of the input sequence provides a single frame view of the sequence and reference junctions (‘Ref. Junctions’) are obtained via extracting the graph structure of the ‘Mean projection’ frame. CC outlines depict the boundaries of the ‘Connected components’ in B. In ‘Ref. Junctions and CC outlines’, Magenta spots show the Ref. Junctions and cyan boundaries show the CC outlines, and the overlay of reference junctions with the CCs shows the extent of movement of junctions within a neighborhood. A single reference junction within a CC is labeled as an Isolated CC (red CC and yellow junctions). Multiple reference junctions within a CC are labeled as Overlapping CC (blue CC and white junctions). Scale bar: 3 *µ*m

#### 2.2.4 Classification of Junction Regions

In our analysis, we utilize the junction projection frame and apply the connected components algorithm to identify regions associated with the movement per junction, referred to as CCs. However, in certain regions of the ER structure, tubules may come into close spatial proximity or overlap with each other, as illustrated in Extended Data Figure 5. In the former case, we can retain individual CCs for the corresponding junctions (Extended Data Figure 5B). For the latter case, we obtain an ‘Overlapping’ CC that encompasses the collective movement (spread) of multiple junctions (Extended Data Figure 5C).

The mean projection frame (Figure 4C) provides a global view of the underlying ER network while accounting for the variable intensity distribution. Using this mean projection frame as input, we extract its graph representation and corresponding junctions within the stable sequence, termed ‘Reference junctions’ (Figure 4C).

To further analyze the connected components (CC) obtained using the connected components algorithm (Figure 4B), we overlay the CC outlines with the reference junctions, as depicted in ‘Ref. Junctions and CC Outlines’ panel of Figure 4C. This visual representation establishes a boundary for the movement of junctions along with their corresponding reference junctions. Leveraging this combined view, we categorize the CCs into the following categories, as explained in the ‘Junctions Distribution’ panel in Figure 4C:

- Isolated CC: A CC with only one associated reference junction.
- Overlapping CC: A CC that contains multiple reference junctions.

### 2.3 Atlastin induces dense tubular networks

Initially, we differentiated Isolated junctions from denser regions of Overlapping junctions to assess the role of ER shaping proteins on tripartite junction formation between ER tubules. The number of reference junctions in Isolated CC regions show consistency across conditions, with only Control and Reticulon 4 conditions showing a significant albeit relatively small difference (Figure 5B), indicating comparable ROIs across conditions.

**Fig. 5.**
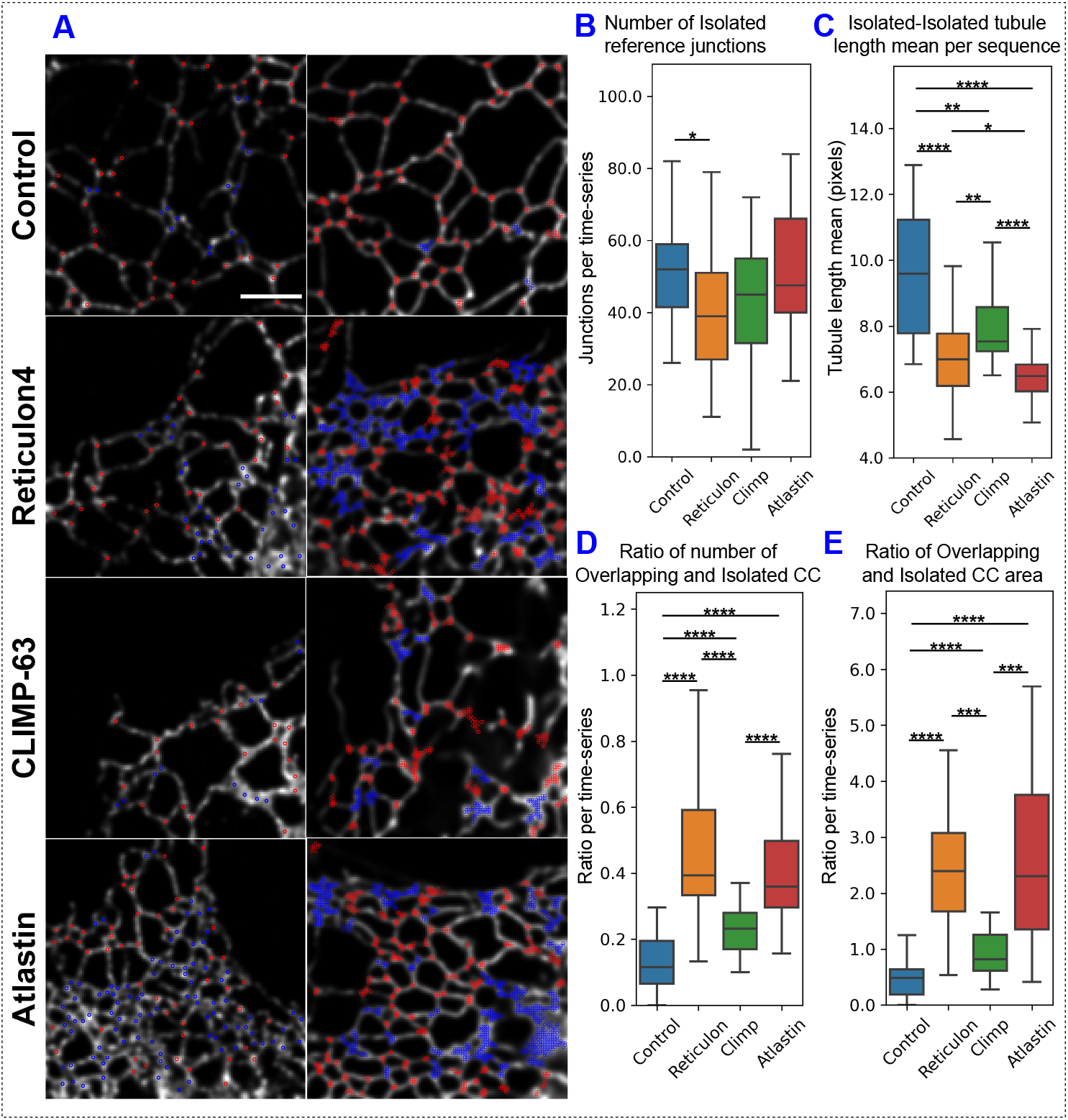
Structural analysis of Isolated and Overlapping CC areas in peripheral ROIs of COS-7 cells. A) Representative images of COS-7 cells transfected with different ER shaping proteins as indicated. ‘Reference junctions’ are labeled as Isolated (Red) or Overlapping (Blue) in column 1 per row. Column two in each row shows the CCs labeled as Isolated (Red) and Overlapping (Blue). B) Number of Isolated reference junctions across conditions, *N* (Control) = 1638, *N* (Reticulon) = 1207, *N* (Climp) = 1371, *N* (Atlastin) = 1292. C) Variation in the mean of tubule length connecting two Isolated junctions per time series. D) Ratio of the number of Overlapping and Isolated areas per time series across conditions. E) Ratio of Overlapping and Isolated CC area per time series across conditions. For panel C), D) and E), *N* (Control) = 31, *N* (Reticulon) = 29, *N* (Climp) = 31, *N* (Atlastin) = 26. Scale bar: 3 *µ*m

To explore the correlation between tubule length and stable junction formation in the ER, we computed the total tubule length per sequence for Control, Reticulon 4, CLIMP-63, and Atlastin transfected cells, based on the mean projection frame (Figure 4C). Tubule length is measured as the distance along the ER skeleton, obtained from the graph representation of the input, where the edges in the graph represent the tubules. Focusing on tubules connecting Isolated CC reference junctions (depicted as ‘Isolated-Isolated tubules’), our analysis reveals that Atlastin induces the shortest tubule length in the ER, followed by Reticulon 4 with slightly longer tubule length (Figure 5C). Statistical analysis based on tubule length demonstrates significant differences between Atlastin and CLIMP-63, as well as Atlastin and Reticulon 4 pairs, evident from the mean plot of tubule length per time-series (Figure 5C).

### 2.4 ER shaping proteins regulated junction dynamics

To investigate junction dynamics, we analyzed the total movement of the junctions based on connected components (CCs) in the projection frame (Figure 4C). In Figure 5A, the first column of each row illustrates the reference junctions labeled as ‘Isolated’ (red spots) and ‘Overlapping’ (blue spots) per group. The second column of each row shows the total movement of Isolated CC (red) and Overlapping CC (blue) per condition.

To quantify the relationship between stable and dynamic regions within the input, we examined the ratio of number of Overlapping and Isolated CCs (Figure 5D as well as the ratio of areas for Overlapping and Isolated CCs (Figure 5E). Reticulon 4 and Atlastin exhibit a significantly increased ratio of Overlapping CCs to Isolated CCs both in terms of junction number and area, compared to Control and CLIMP-63 expressing cells. CLIMP-63 displays a significantly lower ratio of junction number and area, indicating reduced junction dynamics but an increase over the Control condition, which shows the highest stability across all conditions. These results suggest that the expression of Atlastin and Reticulon 4 promotes dynamic, overlapping interactions of ER junctions to a significantly larger extent than CLIMP-63 expression, which resembles the Control.

The extension of the analysis to HeLa cells showed that, consistent with COS-7 cells, Reticulon 4 and Atlastin induced an increase in the number and area of Overlapping CCs relative to Isolated CCs (Figure 6). However, CLIMP-63 increased the Overlapping to Isolated CC ratio to a similar extent as Reticulon and Atlastin which was not observed in COS-7 cells. We interpret this to reflect the increased spreading of COS-7 relative to HeLa cells such that the selected peripheral ROIs include more sheets in HeLa cells than in COS-7 cells.

**Fig. 6.**
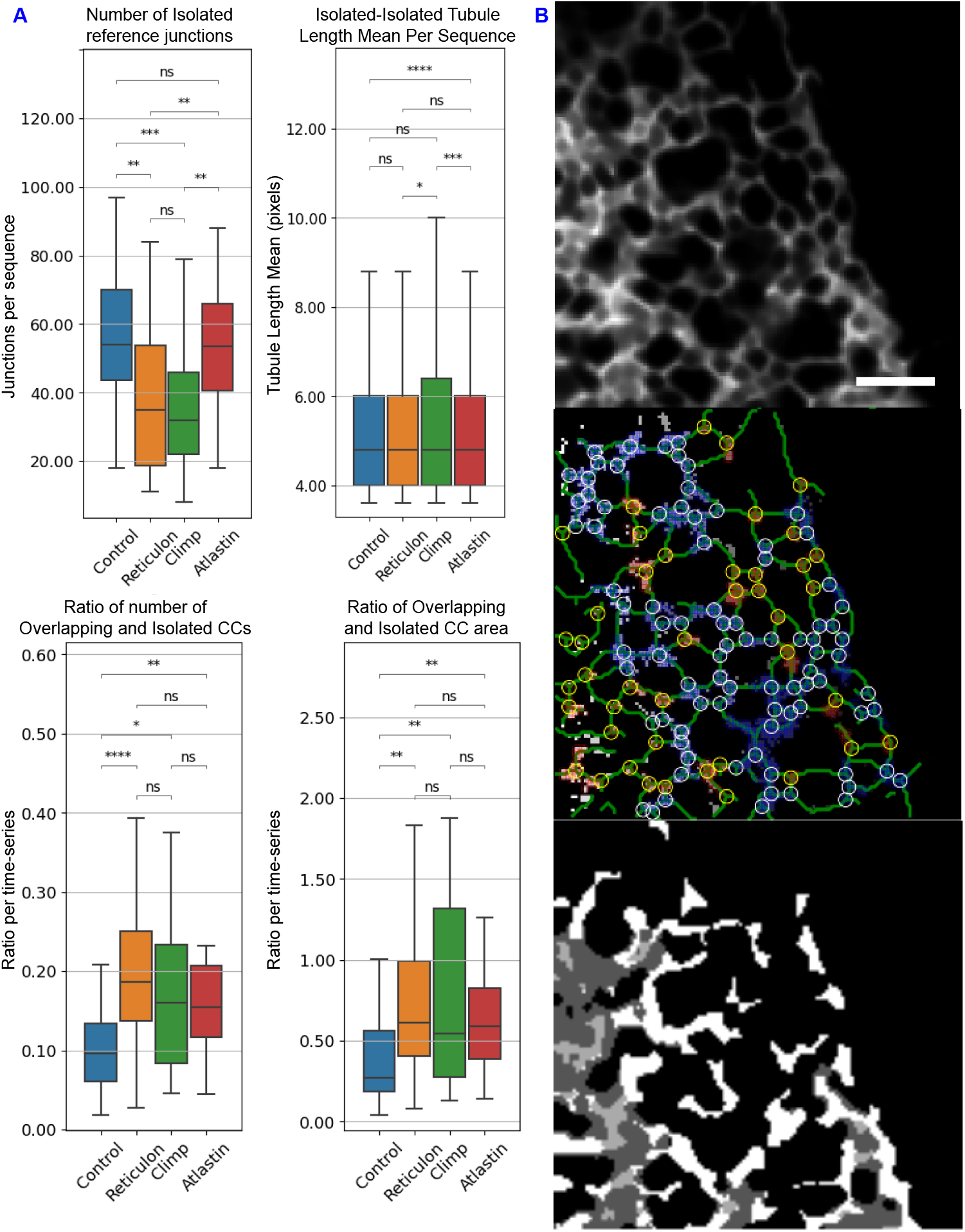
Structural analysis of Isolated and Overlapping CC areas in peripheral ROIs of HeLa cells. A) For HeLa cells transfected with different ER shaping proteins, the following outputs are presented: Number of Isolated reference junctions across conditions; Variation in the mean of tubule length connecting two Isolated junctions per time series; Ratio of the number of Overlapping and Isolated areas per time series across conditions; Ratio of Overlapping and Isolated CC area per time series across conditions. *N* (Control) = 39, *N* (Reticulon) = 28, *N* (Climp) = 25, *N* (Atlastin) = 27. B) Representative image of peripheral ROI of CLIMP-63 transfected HeLa cells. nERdy output, middle panel, shows reference junctions of overlapping CCs as blue with with white rings and reference junctions of isolated CCs as red with yellow rings. Many sheet-like structures (bottom panel) are identified as overlapping CCs. Scale bar: 3 *µ*m

### 2.5 ER shaping proteins regulated tubular matrix dynamics

To validate that the Overlapping CCs induced by Reticulon and Atlastin are due to the enhanced junction dynamics, we evaluated individual CC area as a measure of junction dynamics. In Figure 7A, both Reticulon 4 and Atlastin transfected COS-7 cells exhibited a significant increase in the area of individual Isolated and Overlapping CCs compared to CLIMP-63 transfected cells and Controls. This increase in Isolated CC area suggests that Reticulon 4 and Atlastin promote the dynamics of individual junctions. However, in the Overlapping CCs of the confocal datasets, the skeleton-image correspondence is noticeably reduced (Figure 7B). To address this, we analyzed peripheral regions of interest (ROIs) from Control, mCherry-Reticulon 4, and mCherry-CLIMP-63 transfected COS-7 cells obtained via 2D STED super-resolution microscopy [13]. Application of nERdy+ to these STED datasets effectively reconstructed the ER network (Extended Data Figure 4), with high skeleton-image correspondence observed in the Overlapping CCs (Figure 7B). As observed for analysis of the confocal data, transfection with Reticulon 4 increased both Isolated and Overlapping CC areas relative to CLIMP-63 and Control conditions (Figure 7A). Furthermore, ER tubule dynamics within the Overlapping CC can be more clearly seen in the STED time-lapse series compared to the confocal time series, revealing that the Overlapping CC of Reticulon 4-transfected COS-7 cells define tubular matrix regions consisting of dynamic ER tubules (Figure 7C, see also Supplemental Data videos).

**Fig. 7.**
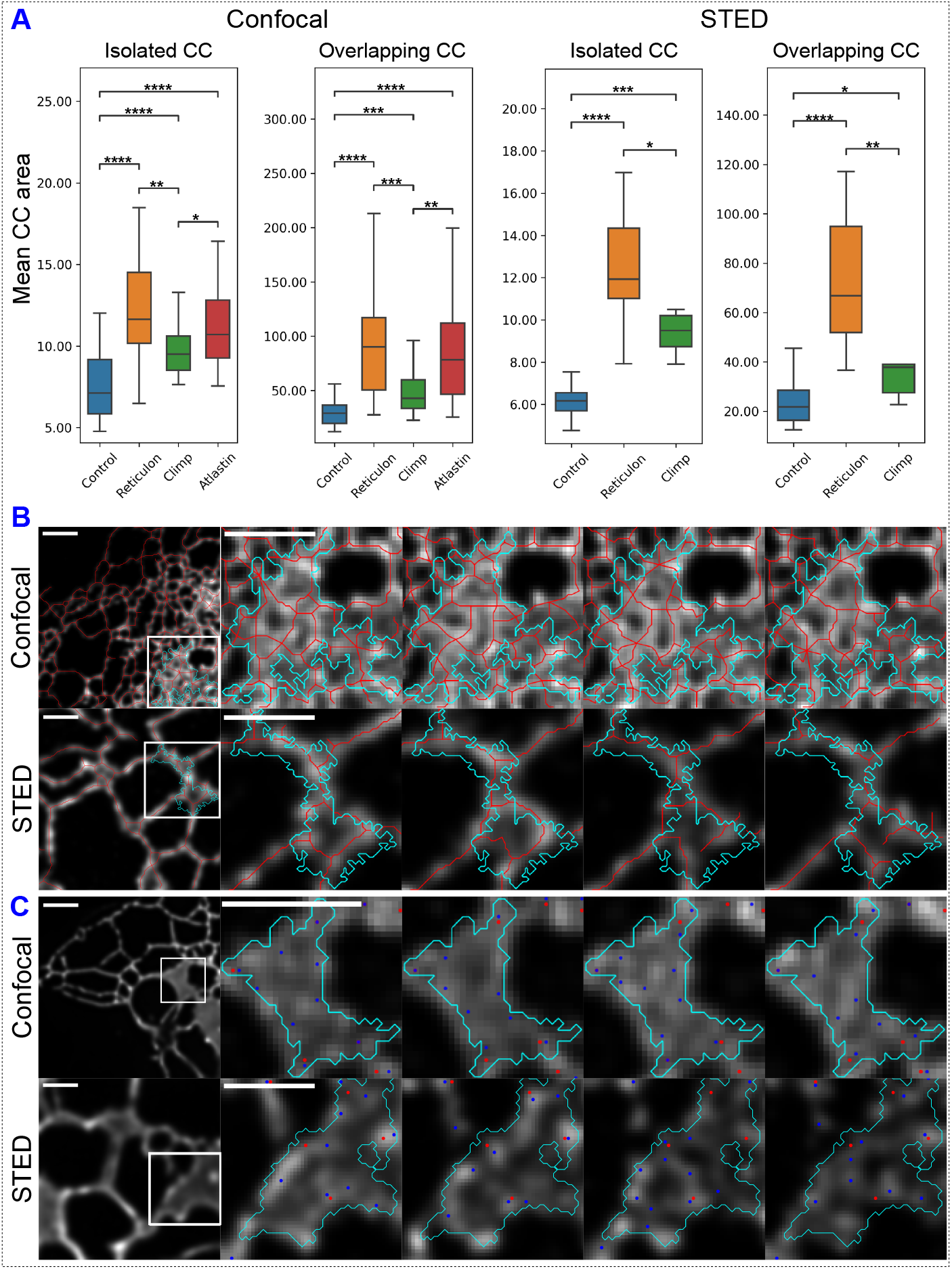
Analysis of junction dynamics. A) Mean CC area is shown per time series, *N* (Control) = 31, *N* (Reticulon) = 29, *N* (Climp) = 31, *N* (Atlastin) = 26. For both confocal and STED data, we observe the highest CC area for Reticulon 4 suggesting more dynamic regions. In confocal, Atlastin follows Reticulon 4 showing high movement. CLIMP-63 shows low movement for junctions whereas Control shows the least movement across the four conditions. These results are consistent for both confocal and STED data. B) Correspondence of skeleton and image across consecutive time-frames for Overlapping CC in confocal and STED data. Scale bar: 3 *µ*m (Confocal), 0.75 *µ*m (STED). C) Mean projections of peripheral ROIs from confocal (CLIMP-63 expressing cell) and STED (Reticulon 4 expressing cell) time series in the leftmost image are shown adjacent to sequential images at a frame interval of 5 frames each (t=[0, 5, 10, 15]) for a tubular matrix region within the ROI (inset, box). Dynamic movement of the tubule and associated junctions (blue) can be more clearly visualized in the Overlapping CC of the STED time series relative to the confocal series. Reference junctions are shown in red. Videos of the presented time series can be seen in Supplemental Data videos. Scale bar: 3 *µ*m (Confocal), 0.75 *µ*m (STED).

## 3 Discussion

The ER shaping proteins CLIMP-63 and Reticulon 4 were originally identified to promote the formation of peripheral ER tubules and sheets, respectively [2, 3]. The tubule and sheet-forming ability of these proteins was based on confocal microscopy analysis of the peripheral ER, and subsequent analysis using high-speed super-resolution microscopy showed that some peripheral sheets were actually dynamic tubular matrices [15]. Here, we develop novel approaches to define the ER network and focus on junction dynamics to distinguish Isolated peripheral tubular regions from more dynamic tubular matrices. We show that tubular matrix regions are enhanced by the Reticulon 4 and Atlastin ER shaping proteins and use super-resolution microscopy to show that they are composed of dynamic tubules.

To segment the ER structure, we employ either a set of image processing steps (nERdy) or train a model (nERdy+). Subsequently, we skeletonize the segmentation output to derive a single-pixel-wide network representation of the ER. Our ground truth includes manually drawn skeletons for the input ER frames, which we dilate to create a ‘GT mask’ for training in the segmentation task. This shift improves contextual understanding of the underlying ER structure, therefore avoiding potential nuances of error in single-pixel-wide structure prediction. This refined GT mask faithfully captures the overall ER structure.

Our first approach, nERdy, utilizes classical image processing techniques. It is primarily motivated by AnalyzER [28] and addresses limitations seen in AnalyzER which involves extensive parameter tuning at each step in their pipeline, particularly problematic for larger experiments. The default parameters in AnalyzER’s GUI-based software do not faithfully represent ER networks in our data (Figure 3C). nERdy reduces the number of adjustable parameters, offering improved performance on our test dataset compared to AnalyzER.

nERdy involves a series of image processing steps designed to extract the tubular ER structure from time-lapse sequences. While we have demonstrated the robustness of our parameter configuration across our samples, we recognize that different imaging modalities may require parameter adjustments. The intensity normalization and histogram equalization steps are essential for enhancing contrast and standardizing intensity levels across different frames and samples. For datasets with varying illumination conditions, we suggest adjusting the normalization range to match the intensity distribution of the new data. Histogram equalization parameters might need to be fine-tuned to avoid over-enhancement of noise. A good practice is to visually inspect a few frames and adjust the parameters accordingly. The morphological operations such as area opening, erosion, and local thresholding help remove noise and fine structures while preserving tubular integrity. In area opening operation, we suggest adjusting the minimum area threshold based on the size of the smallest tubular structures to be preserved. For larger structures, the threshold can be increased to filter out smaller noise components. In erosion operation, the structuring element size and shape can be modified based on the thickness of the tubular structures. For thinner tubes, we suggest using smaller structuring elements. In local thresholding, the parameters should be set considering the local contrast variations. Adaptive thresholding methods, such as Otsu or Sauvola, might be beneficial for datasets with high variability. The Jerman Enhancement filter enhances tubular structures by emphasizing line-like features. The scale and sensitivity parameters of the Jerman Enhancement Filter should be adjusted according to the resolution and specific characteristics of the tubular structures in the dataset. Higher resolutions might require finer scales, while lower resolutions might need broader scales.

To utilize nERdy on a custom dataset, start with the default parameters provided in our method and apply these to a subset of your data to evaluate the segmentation results. Based on the initial results, iteratively adjust one parameter at a time, documenting the changes and their impacts to systematically approach the optimal configuration. Use visual inspection of segmentation overlays on the original images to validate performance, ensuring that the segmented structures align well with the expected ER morphology. Consider using automated parameter tuning techniques, such as grid search or random search, to explore a wider range of parameter values systematically. For high-resolution confocal microscopy, setting the scale parameters of the Jerman Enhancement Filter to finer values and reducing the structuring element sizes in morphological operations can be helpful. In contrast, for low-resolution widefield microscopy, broader scales for the enhancement filter and larger structuring elements might be more appropriate to capture the coarser details of the ER network.

Our deep learning model, nERdy+, demonstrates superior overall performance across confocal and STED time-lapse series and protein conditions. However, it faces limitations in specific graph measures, particularly noticeable in STED data due to intricate annotations requiring four times the annotation time of confocal data. Given the ease of acquisition and annotation convenience of confocal data, we focused our training on it. Consequently, nERdy+ experiences a drop in performance when applied to STED data due to ‘domain shift’ [39, 40], a common issue in machine learning. Nevertheless, nERdy still performs admirably in both confocal and STED data, showcasing robustness even though its parameters are set based solely on confocal data.

In our study, we employed the Dice score, Jaccard Index, and F1-score to evaluate the performance of our model. Each metric offers distinct advantages and insights. The Dice score is particularly sensitive to the overlap between predicted and actual segments, making it invaluable for tasks requiring precise boundary delineation in the case of ER tubules. The Jaccard Index, or Intersection over Union, provides a balanced measure that is less sensitive to class imbalance, making it suitable for overall accuracy of segmented ER regions. The F1-score, combining precision and recall, is crucial for scenarios where both false positives and false negatives have significant consequences, offering a balanced view especially in the cases where ER structures may vary in density and complexity. By utilizing these metrics together, we ensure a robust and comprehensive assessment of our model’s performance, handling the challenges of ER segmentation.

Deep learning based approaches, ERnet [29] and ERnet-v2 [30], avoid the need to calibrate parameters in image processing. nERdy+ aims to provide an efficient, adaptive, and more robust alternative to these methods. While deep learning based methods typically excel with ample training data and ground truth, obtaining annotations for ER segmentation, particularly concerning medial lines in ER skeletons, remains challenging.

The classical approach to learning representations from limited data involves data augmentation techniques [41], such as rotation, which provides different orientations of the input to share learnable neural network parameters across these variations. Although this augmentation implies a form of equivariance, it may lead to learning only approximate equivariance when the network architecture has insufficient capacity, and thus the invariance learned on the training set may not generalize equally well to a test set [42]. Data augmentation requires the generation of augmented samples, leading to increased memory/storage footprint and compute cycles. In contrast, directly incorporating symmetry information of the data into the network architecture, rather than augmenting the training data, can improve performance [43]. With nERdy+, we’ve introduced a method based on equivariant neural networks, which significantly improves handling limited data by preserving data symmetries.

A promising approach within the limited data context is few-shot learning [44], allowing models to learn from a minimal number of instances per class. Future research may explore synergies between equivariant networks and few-shot learning [45] for ER analysis. However, in the absence of ground truth and training data, nERdy can provide strong baseline performance. Recent approaches such as self-supervised learning [46] aim to alleviate the dependency on large supervised training data and represent a promising avenue for future research in ER segmentation.

The networks extracted across consecutive frames depict junction dynamics over time. A mean projection frame, aggregating all networks, provides a comprehensive snapshot of overall network movement. This frame highlights stable areas with higher intensity and dynamic regions with lower intensity, offering a stable representation of ER movement. However, tracking individual junctions poses challenges due to diverse network representations in each frame and the proximity of junctions leading to overlap in (*X, Y*) pixel locations over time. To address these challenges and study the junction dynamics, we utilize both the stable view from the projection frame and the dynamic view via changing junction locations from subsequent frames. This approach captures nuanced protein dynamics and differences across conditions. The low-movement regions are denoted as ‘Isolated CC’ and high-movement regions are denoted as ‘Overlapping CC’. We observe that Isolated CCs maintain consistent sizes across confocal and STED data, while Overlapping CCs show reduced size in STED data, underscoring the impact of resolution on detecting ER dynamics.

Analysis of the peripheral tubular network, connecting Isolated CCs, reveals that overexpression of Atlastin, and to a lesser extent Reticulon 4, most significantly reduces mean tubule length between Isolated junctions. This is consistent with the established role of these two ER shaping proteins in promoting the formation of peripheral ER tubules and the tripartite junctions that connect them [2, 8, 9]. Of particular interest was the dramatic increase in Overlapping CC regions induced by Atlastin and Reticulon 4. This supports a role for these ER tubule-forming proteins in the establishment not only of the extended peripheral ER network but also of denser tubular matrices. These effects were attributed to increased junction dynamics as the expression of Atlastin and Reticulon 4 increased the area of Isolated CCs, reflective of the movement of peripheral tripartite junctions. Mechanisms underlying ER junction dynamics involve close interaction and interdependence between the ER network and the microtubule cytoskeleton [10, 12, 47, 48]. Our data suggests that Atlastin and Reticulon 4 promote junction dynamics leading to the formation of dense tubular matrices.

While intensity variations are present in the Overlapping CCs observed in confocal time series (Figure 7B), defining these regions as either peripheral sheets or tubular matrices proves challenging with confocal microscopy. Annotating ER tubules in these regions, whether by nERdy or manual annotation, faces significant hurdles; the varied junction distribution may reflect annotation challenges as much as tubule dynamics. Extension of nERdy+ to STED super-resolution time-lapse series, where annotation of ER tubules in individual frames is feasible (Figure 3A), clearly shows that Overlapping CC regions are composed of dynamic tubules. Consistent with our confocal analysis, Reticulon 4 induced the more extensive formation of tubular matrices than CLIMP-63 expression.

CLIMP-63 expression in COS-7 cells induces the extensive formation of ER sheets [3, 13, 14]. In our experiments, CLIMP-63 overexpression led to a modest increase in Overlapping CCs compared to the ability of Reticulon 4 and Atlastin to induce extensive tubular matrices in COS-7 cells. Peripheral sheets identified by confocal microscopy may include both extended sheets as well as tubular matrices that will be difficult to distinguish by nERdy+. The inclusion of some sheets, alongside tubular matrices in the peripheral ROIs studied may be responsible for the small increase in Overlapping CCs relative to Control observed upon CLIMP-63 overexpression. Indeed, this became evident upon analysis of HeLa cells in which CLIMP-63 induced more Overlapping CCs than in COS-7 cells. This is likely due to the increased spreading of COS-7 cells such that the peripheral ROIs studies do not include as many sheets. Dense tubular matrices induced by Atlastin and Reticulon 4 would appear to be intermediate ER structures between the extended peripheral ER networks and more central CLIMP-63-dependent ER sheets [49].

While super-resolution microscopy enables the visualization of dynamic tubular matrices in cultured cells in this and other published studies [15], convoluted tubular smooth ER networks have long been observed by EM in various tissues [49]. The demonstration here that Reticulon 4 and Atlastin promote tubular matrix formation supports a role for these ER shaping proteins in regulating smooth ER network expression in tissues. Together with earlier results showing that ER shaping proteins regulate the dynamics of peripheral ER tubules and ER sheets [13, 14], these results highlight the complex role of ER shaping proteins in the dynamic organization of the ER.

## 4 Methods

### 4.1 Cells and reagents: Plasmids

ERmoxGFP was a gift from Dr. Erik Snapp (Albert Einstein College of Medicine, present Howard Hughes Medical Institute Janelia Research Campus, Virginia) (Addgene plasmid # 68072), mCherry-CLIMP-63 from Dr. Tom Rapoport (Harvard University, Massachusetts), mCherry-RTN4A and mCherry-ATL1 from Addgene (Addgene plasmid #86683 and #86678, respectively).

### 4.2 Cells and reagents: Cell line

HeLa cell line was acquired from ATCC and authenticated by Short Tandem Repeat (STR) profiling at the TCAG Genetic Analysis Facility (Hospital for Sick Kids, Toronto, ON, Canada www.tcag.ca/facilities/geneticAnalysis.html). COS-7 cell line (CLS Cat# 605470/p532_COS-7, RRID:CVCL_0224) was acquired from ATCC and gifted from Ann-Marie Craig (UBC). All cell lines were tested regularly for mycoplasma infection by PCR (ABM, Richmond, BC, Canada).

COS-7 and HeLa cells were grown at 37°C with 5% CO2 in complete Dulbecco’s Modified Eagle’s Medium (DMEM) (Thermo Fisher Scientific, USA) containing 10% FBS (Thermo Fisher Scientific, USA) and 1% L-Glutamine (Thermo Fisher Scientific, USA) unless otherwise stated. Plasmids were transfected in COS-7 cells with Effectene (Qiagen, Germany) according to the manufacturer’s protocols for 22 hours. For live cell imaging, 15K COS-7 or HeLa cells were plated in ibidi 8-well m-slides (cat. No: 80827) with #1.5H (170 µm *±* 5 µm) D 263 M Schott glass and incubated for 24 hours. The next day, cells were transiently transfected with ERmox-GFP alone or with mCherry-Climp63, -Reticulon-4A or -Atlastin using Lipofectamine-2000 (Invitrogen, USA) following the manufacturer’s protocol and allowed to grow for an additional 24 hours in the incubator. Before imaging, complete DMEM medium was replaced with DMEM live-cell imaging medium (Sigma, USA) without sodium bicarbonate and phenol red, supplemented with 1% L-glutamine, 10% FBS, and 10% HEPES (Thermo Fisher Scientific, USA). Imaging was conducted at 37^°^C.

### 4.3 Image acquisition

Confocal and gSTED imaging were performed with the 100X/1.4 Oil HC PL APO CS2 objective of a Leica TCS SP8 3X STED microscope (Leica, Germany) equipped with a white light laser, HyD detectors, and Leica Application Suite X (LAS X) software. GFP was excited at 488 nm (for confocal or STED) and depleted using the 592 nm depletion laser (STED only). mCherry was excited at 584 nm. Both confocal and STED images were deconvolved using Huygens Professional software (Scientific Volume Imaging, The Netherlands). Time-gated fluorescence detection was used for STED to further improve lateral resolution;; estimated XY FWHM values obtained from the theoretical PSFs for STED GFP images were 78 nm for 2D live analysis. Live cell time-lapse imaging of GFP was conducted on select square ROIs in the periphery of the cell. The STED time-lapse series dataset has been previously published [13].

### 4.4 Implementation details: nERdy

Network analysis of Endoplasmic Reticulum dynamics (nERdy) combines various morphological operations. To begin, the input sample is normalized within the range of zero to one, followed by Contrast Limited Adaptive Histogram Equalization (CLAHE; [50]). CLAHE helps in improving the local area contrast within the image. Next, the morphological area opening operation is performed with an area threshold value set to two. This operation removes very small objects and restores the remaining objects to their original size. Subsequently, an erosion operation is carried out using a 3 *×* 3 square structuring element with a connectivity of one. The center pixel of the underlying input is preserved if all the pixels in the 3 *×* 3 neighborhood belong to the foreground region. This erosion step brings us closer to a thin version of the input tubular structure. As a last step, local thresholding is performed using a block size of 3. Local thresholding enhances the structure in the input while adaptively removing very low signal values. The Jerman Enhancement [34] method analyzes the local intensity structure using the Hessian matrix of the input. The relative eigenvalues of the Hessian matrix are used to identify the regions with high vessel-like or tubular structures.

The morphological operations in nERdy and junction analysis routines were developed using Python and the image processing library, scikit-image [51]. Only in the case of Jerman Enhancement Filter [34], we use the open source MATLAB code available at https://github.com/timjerman/JermanEnhancementFilter and use the MATLAB Engine API in python to integrate the tubular enhancement routine in our nERdy pipeline.

### 4.5 Implementation details: Equivariance

Equivariance is a property of a mathematical operation whereby its output changes predictably in response to the transformations applied to its input. For instance, rotating an image of a dog by 90 degrees should rotate the dog segmentation mask by 90 degrees as well. Considering an input x in *R*^2^ space (e.g., an *XY* coordinate), formally, for a transformation *T* applied to the input *x*, and a function *f* representing the neural network, equivariance can be expressed as:

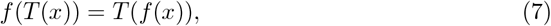

where *T* (*x*) is the transformed input, *f* (*x*) is the network’s output for the original input and *T* (*f* (*x*)) is the transformed output. Another property of interest is invariance, formally denoted as:

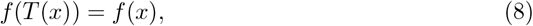

where the transformation does not affect the output of the neural network. For example, the class predicted for an image of a dog should not change to another class when the image is rotated (the image remains an image of a dog). We illustrate the transformation for a single ER frame in Extended Data Figure 2.

### 4.6 Implementation details: Dihedral group and G-convolutions

The traditional data augmentation approach encourages the model to learn symmetries incurred by augmentation, but it does not ensure equivariance. In a standard convolutional neural network (CNN), each convolutional layer consists of a set of learnable kernels. Here, we adopt an equivariant approach relying on ideas from group theory, specifically dihedral group *D*_*n*_, to ensure that the model utilizes the symmetries during training. A dihedral group is defined as the set of symmetries preserving the shape of a regular polygon with n sides. Comprising rotations and reflections, *D*_*n*_ captures the interplay between angular rotations and mirror symmetries intrinsic to these polygons. The group has a total of 2*n* elements, corresponding to *n* rotations and *n* reflections.

In contrast to CNNs, the equivariant neural networks utilize group convolutions, also known as G-convolutions, to learn equivariant feature maps [52]. Given a feature map *f*, a group element *g* and *K*_*g*_ as the kernel associated with the group element *g*, the group convolution operation is defined as:

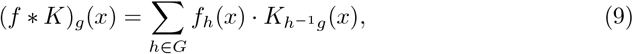

where *f*_*h*_(*x*) represents the feature map *f* transformed by the group element *h*, 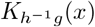 represents the kernel *K* transformed by the composition of group elements *h*^−1^ and *g, x* denotes the spatial coordinates of the feature map.

With G-convolutions, the learnable kernels transform through the action of a group *G*. Considering the *D*4 group in our experiments, the action involves rotation and reflection. Consequently, instead of having *k* individual kernels as in a convolutional layer, the equivariant layer now encompasses |*G*|× *k* kernels, where |*G*| denotes the order of the group. Each input is convolved with a set of kernels corresponding to each transformation in the *D*4 group. This ensures that the network captures consistent features across different orientations of the input. Subsequently, after convolution, the feature maps are transformed back to the original orientation, aligning them with the input. This process builds a single equivariant layer and continues in deeper layers by considering input feature maps rather than raw data. This approach enables the network to capture complex patterns while preserving equivariance to the transformations.

### 4.7 Implementation details: nERdy+

We build the equivariant layers as suggested in [52] using the PyTorch framework. Our architecture consists of six equivariant layers followed by a transposed convolution layer (Extended Data Table 1). Following [52], we apply the group spatial max pooling operation only once after the second equivariant convolution layer. This operation is applied over each feature map. Following the last convolution layer (layer number 7), the feature maps are concatenated along the channel dimension. This output is passed to the transposed convolution operation which provides the probability map. nERdy+ has a total of 1,549,953 trainable parameters. We train nERdy+ using the binary cross entropy loss function given as:

**Table 1.** Architecture for nERdy+. The P4MConvZ2 layer takes input and provides the P4M transformation (denoting four rotations and mirroring/ reflection). P4MConvP4M layer takes the transformed input and provides transformed output, where the transformations are ruled by the specified group (D4 in this case). Group Spatial Max Pooling performs pooling per feature map. ConvTranspose2d layer applies transposed convolution operation over the transformed input from the previous layer to provide the final segmentation output.

| Layer | Output shape | Parameters |
| --- | --- | --- |
| Input | [1, 128, 128] | 0 |
| P4MConvZ2 | [-1, 32, 8, 128, 128] | 320 |
| P4MConvP4M | [-1, 32, 8, 128, 128] | 73,760 |
| Group Spatial Max pooling | [-1, 32, 8, 64, 64] | 0 |
| P4MConvP4M | [-1, 64, 8, 64, 64] | 147,520 |
| P4MConvP4M | [-1, 128, 8, 64, 64] | 589,952 |
| P4MConvP4M | [-1, 64, 8, 64, 64] | 589,888 |
| P4MConvP4M | [-1, 32, 8, 64, 64] | 147,488 |
| ConvTranspose2d | [-1, 1, 128, 128] | 1,025 |

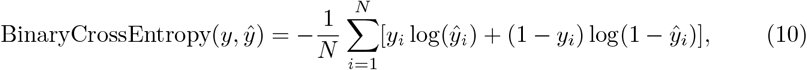

where *y* represents the ground truth labels (‘GT mask’; Figure 2), ŷ represents the predicted probabilities (‘Predicted segmentation mask’; Figure 2), *N* is the total number of samples (validation samples in confocal and STED data), and *y*_*i*_, ŷ_*i*_ are the *i*^*th*^ elements of *y*, ŷ, respectively. The transposed convolution operation provides output as logits (raw model output) instead of probabilities and thus we apply the sigmoid activation function over the prediction logits to obtain probabilities (implemented as BCEWithLogitsLoss in PyTorch). The weights of the model are updated using VectorAdam optimizer [53]. Adam [54] is a widely used optimizer in machine learning tasks, but it provides per-coordinate moment updates. VectorAdam strives for rotation equivariance via considering the vector-valued structure of the model parameters. In our experiments, the learning rate for VectorAdam is set to 8e− 4, *β*_1_, which controls the weight decay for first moment estimates, is set to 0.9, and *β*_2_, which controls the weight decay for second moment estimates, is set to 0.999. In addition, an epsilon parameter acts as a small positive constant added to the second moment estimates to avoid division by zero. We set epsilon as 1e-8. We utilize NVidia GeForce GTX 1080 Ti GPU with 12GB of RAM for training. The batch size is set to 32 and the model is trained for 100 epochs. The model takes ∼1 second per epoch leading to a total time of 1.67 minutes for training. The segmentation evaluation measures are implemented in Python.

### 4.8 Implementation details: Skeleton to Graph

A skeleton, here, is a single pixel-wide binary input. Initially, each pixel in the skeleton is considered a node in the graph, and the 3-way bifurcation locations are identified based on the degree (number of immediate connections) of each node. This results in a new representation of the graph, where nodes with a degree of 3 or higher act as the new nodes, and the pixels or nodes from the initial graph representation serve as the edge coordinates connecting the new higher-degree nodes. The tubular connections between these nodes form the edges of the graph, representing the tubules in the structure. For skeleton-to-graph conversion, we use the sknw module from ImagePy library [55] and available at https://github.com/Image-Py/sknw. Similar to ERnet-v2 [30], the extracted ER structure graphs are undirected and unweighted. Graph measures are obtained using NetworkX package [56].

### 4.9 Statistics and Reproducibility

Statistical analyses were performed using a two-sided Mann–Whitney U test with Bonferroni correction. P-values were reported as follows: ns: *p >* 0.05;∗: *p* ≤ 0.05;∗∗: *p* ≤ 0.01;∗∗∗: *p* ≤ 0.001;∗∗∗∗: *p* ≤ 0.0001. Sample sizes (*N*) for each analysis are provided in the figure legends. All experiments were conducted in three biological replicates, with each replicate comprising at least seven independent cell regions. Replicates were defined as independent cells or fields of view acquired under identical experimental conditions. All experiments were reproducible across biological replicates.

## 5 Data availability

All datasets used to develop and test nERdy, nERdy+, and the findings presented in this manuscript are publicly available at the figshare repository [57].

## 6 Code availability

The code for both nERdy and nERdy+ is available on GitHub at https://github.com/-NanoscopyAI/nERdy.

## 7 Author contributions

A.S. performed the data analysis, developed the computational pipeline for nERdy, nERdy+ and wrote the article. G.G. conducted the experiments, acquired the images, and edited the article. B.C. gave advice and edited the article. B.J. conducted the experiments. I.R.N. and G.H. conceptualized the study, supervised the research, and wrote the article.

## 9 Extended Data

**Extended Data Figure 1.**
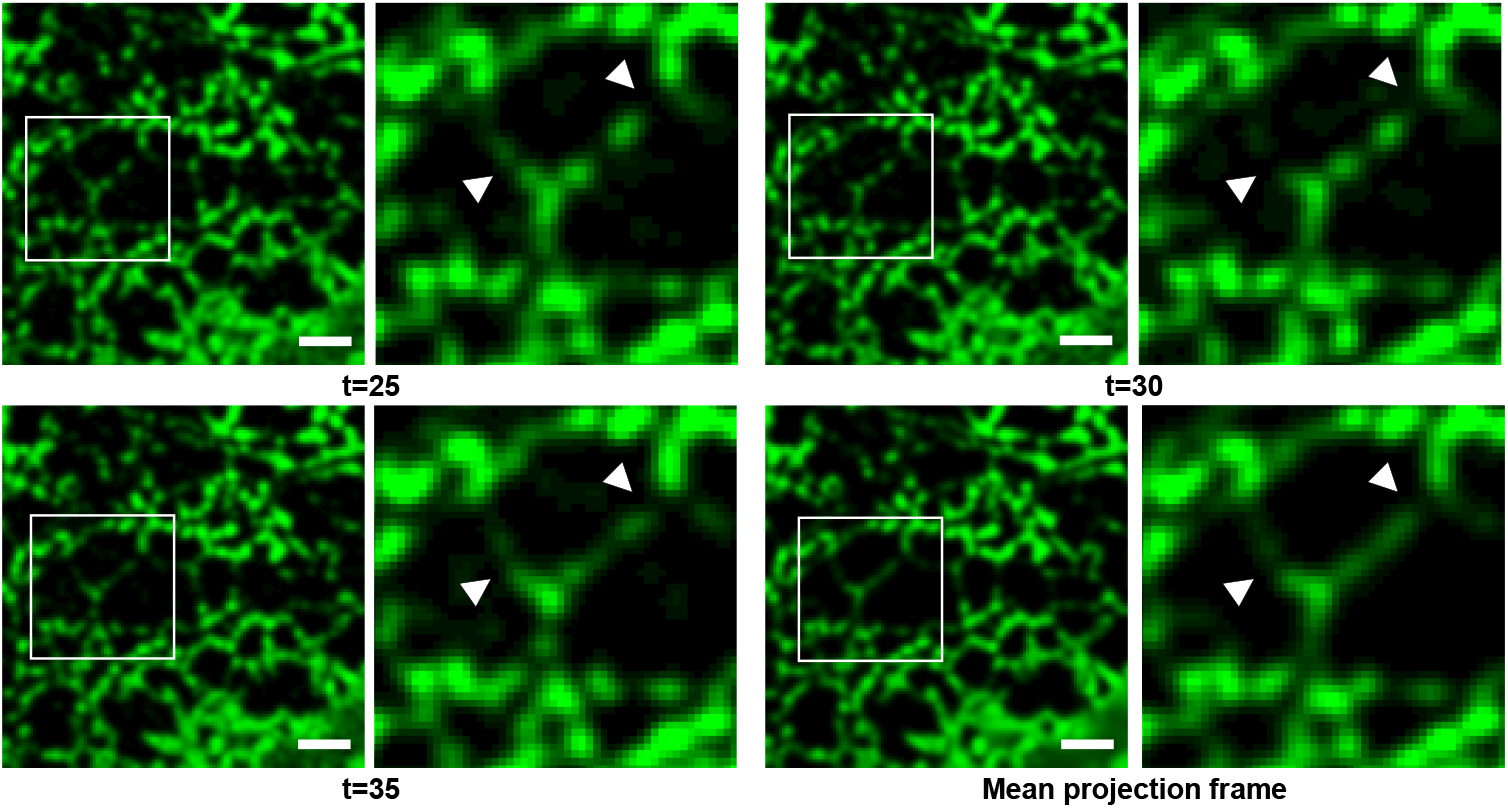
ER network discontinuities in confocal data. For an ROI of COS7 cell transfected with ERmoxGFP/ Atlastin, we observe a decrease in the signal (deformation) and an increase in the signal (formation) representing the tubules at specified locations across three distinct time steps. The ER structure is not continuous in individual frames but shows an intact nature in the mean projection frame as the projection frame compensates for the near-zero intensity regions by averaging over all frames. Scale bar: 3 *µ*m

**Extended Data Figure 2.**
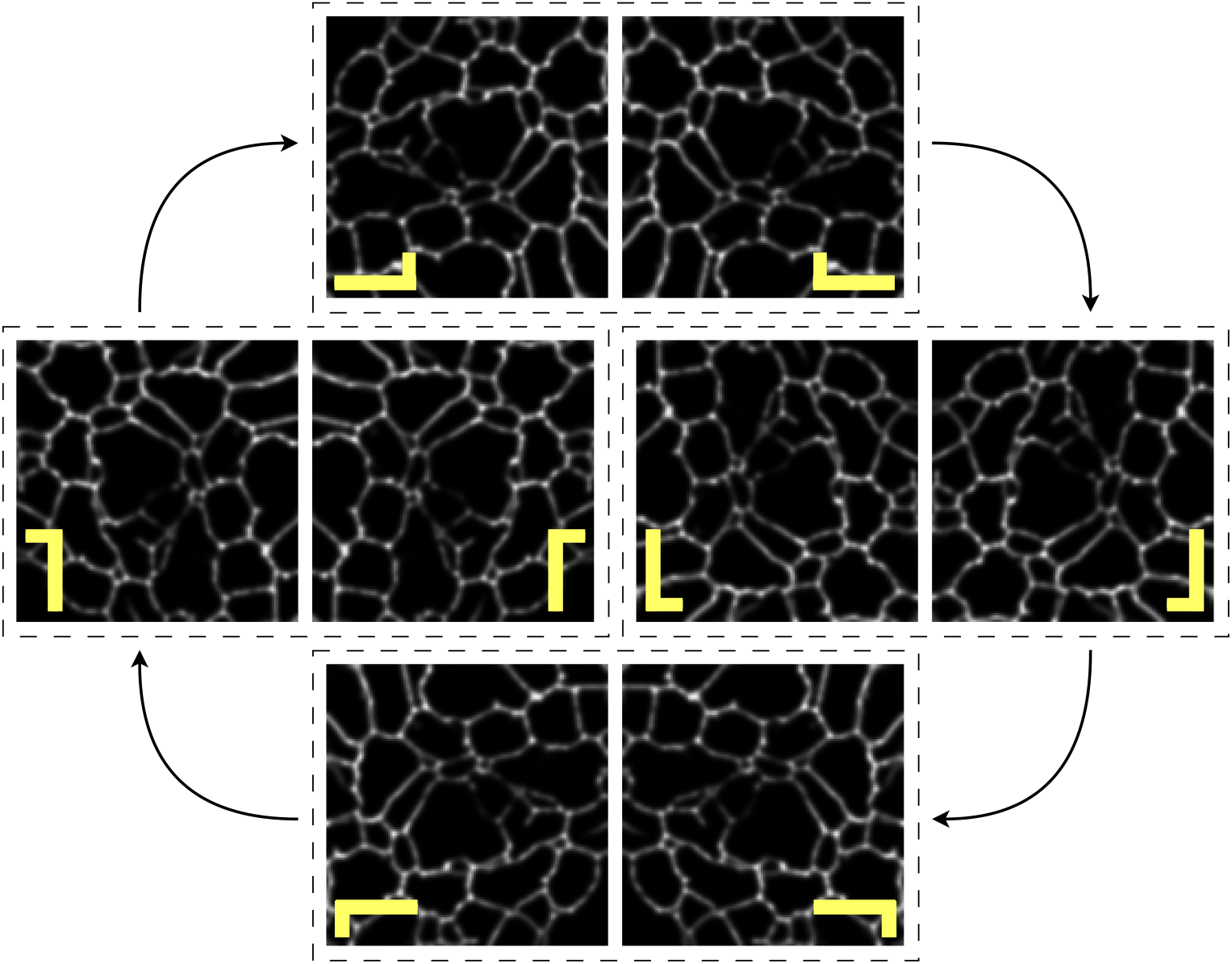
Illustration of the D4 group. The elements of the D4 group each are rotated by π*/*2 radians and mirrored around x=0, resulting in a group order |*D*4| = 8. Example of group action applied to a single ER frame. We add an L-shaped glyph in yellow to clarify the different transformations applied over the input.

**Extended Data Figure 3.**
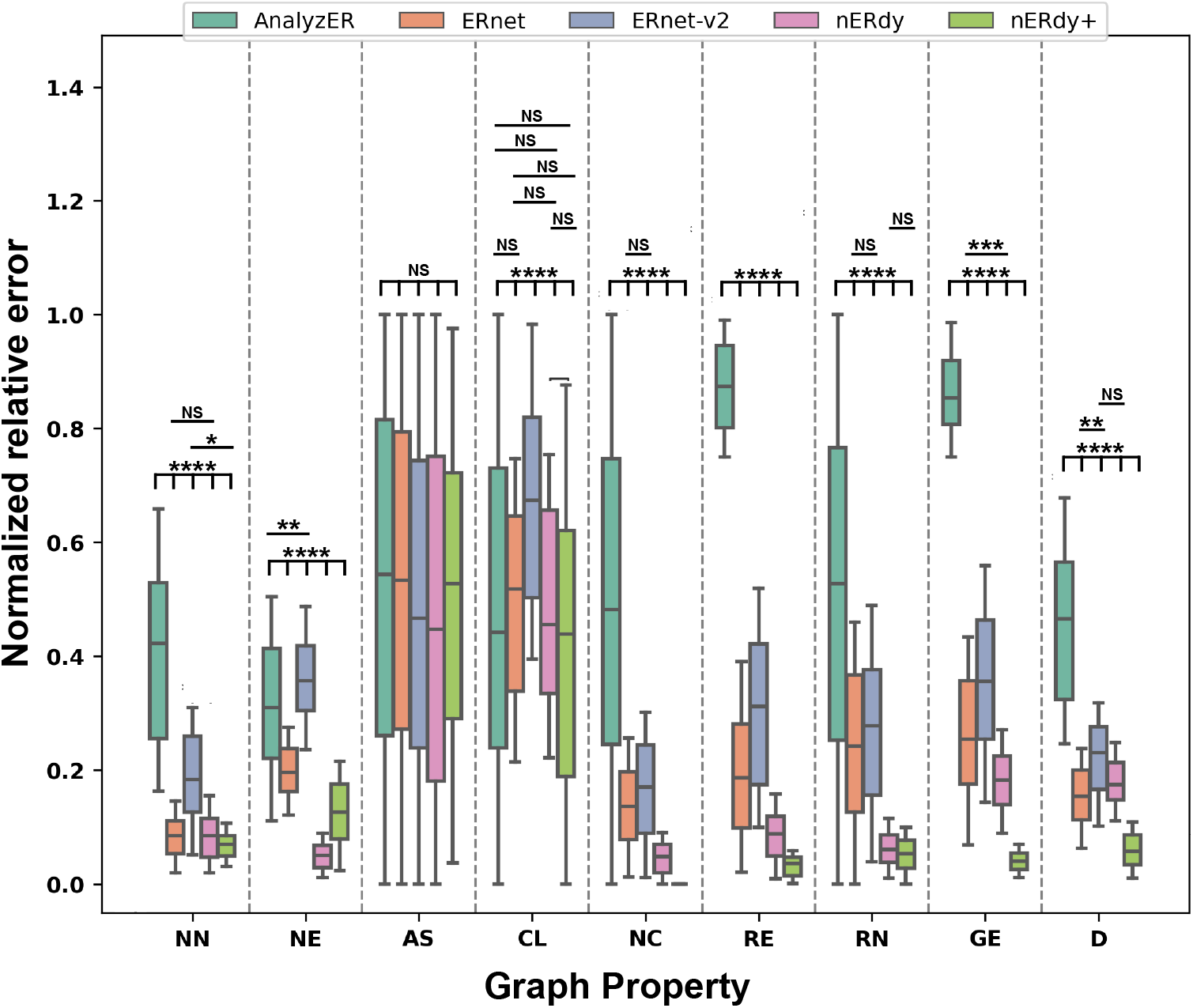
Quantitative evaluation of error in graph reconstruction across the methods in confocal data (*N* = 21). The graphs are constructed using the segmentation output from each method. For each method, we compare the output graph with the ground truth graph using various graph properties and calculate the relative error. The error is normalized in 0-1. The properties include: the number of nodes in the graph (NN), number of edges in the graph (NE), assortativity coefficient of the graph (AS), clustering coefficient of the graph (CL), Number of connected components in the graph (NC), ratio of number of nodes in the biggest subgraph and the number of nodes in the graph (RN), ratio of number of edges in the biggest subgraph and the number of edges in the graph (RE), global efficiency of the graph (GE) and density of the graph (D). For confocal time-lapse series data, nERdy+ shows the smallest relative error for all the graph properties, whereas AnalyzER shows the highest error across all graph properties. The wide confidence intervals in AnalyzER suggest inconsistent performance across data samples and thus a lack of adaptability. ERnet and nERdy show close performance on the majority of the properties with ERnet-v2 showing better performance in properties such as density, and global efficiency.

**Extended Data Figure 4.**
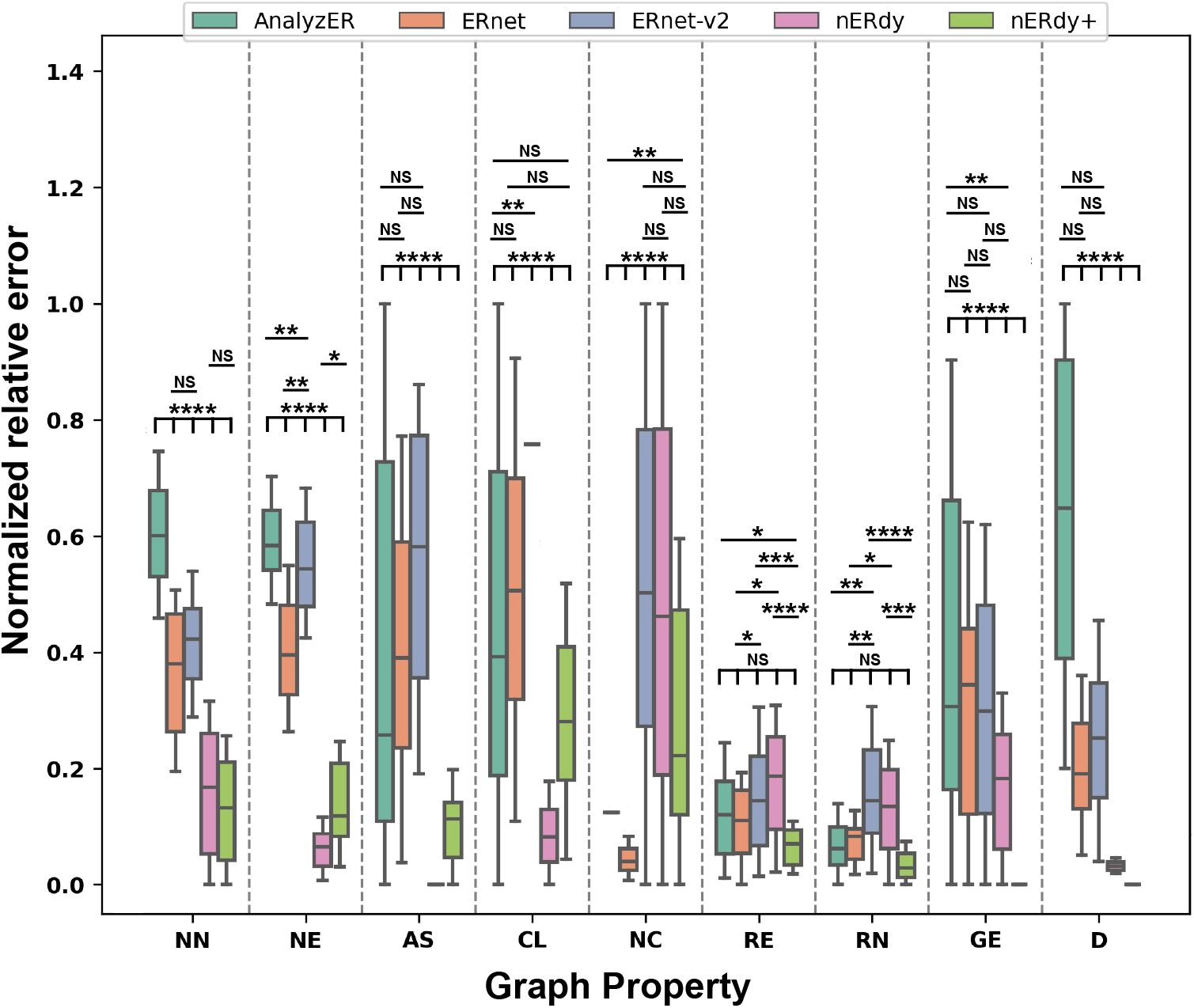
Quantitative evaluation of error in graph reconstruction in STED data (*N* = 35). nERdy shows the least relative error in 4 properties: number of nodes (NN), number of edges (NE), assortativity coefficient (AS), and clustering coefficient (CL). ERnet-v2 shows the least relative error in the number of components (NC), and global efficiency (GE). ERnet also shows the least relative error in 2 metrics, namely, the ratio of edges (RE) and the ratio of nodes (RN).

**Extended Data Figure 5.**
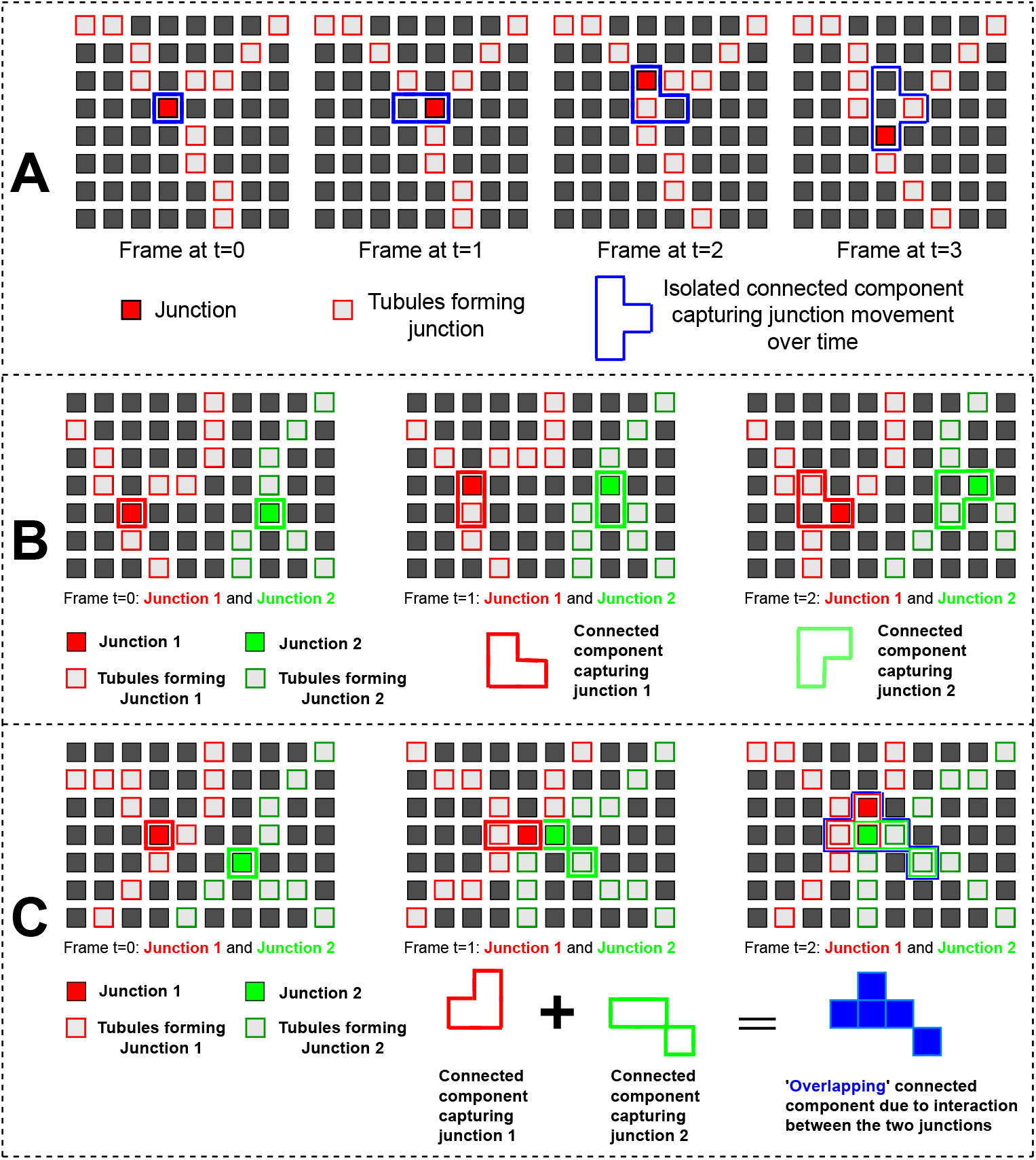
Schematic of tubule movement and CC formation analysis. A) Movement of tubules and junctions leads to the formation of junction movement region depicted as CC. At each subsequent step, the new junction location (denoted in red) is incorporated in the connected component (denoted using blue boundary). The total movement of a junction within a time series is captured in the CC. B) We observe the movement of two individual tubules without any interaction between them. These tubules form the Isolated CCs. C) Two spatially close tubules and their interaction over time leading to Overlapping CCs.

## References

[1] Westrate, L.M., Lee, J.E., Prinz, W.A., Voeltz, G.K.: Form follows function: the importance of endoplasmic reticulum shape. Annu Rev Biochem 84, 791–811 (2015) 10.1146/annurev-biochem-072711-163501

[2] Voeltz, G.K., Prinz, W.A., Shibata, Y., Rist, J.M., Rapoport, T.A.: A class of membrane proteins shaping the tubular endoplasmic reticulum. Cell 124(3), 573–586 (2006)

[3] Shibata, Y., Shemesh, T., Prinz, W.A., Palazzo, A.F., Kozlov, M.M., Rapoport, T.A.: Mechanisms determining the morphology of the peripheral er. Cell 143(5), 774–788 (2010)

[4] Wang, B., Zhao, Z., Xiong, M., Yan, R., Xu, K.: The endoplasmic reticulum adopts two distinct tubule forms. Proceedings of the National Academy of Sciences 119(18), 2117559119 (2022)

[5] Tolley, N., Sparkes, I.A., Hunter, P.R., Craddock, C.P., Nuttall, J., Roberts, L.M., Hawes, C., Pedrazzini, E., Frigerio, L.: Overexpression of a plant reticulon remodels the lumen of the cortical endoplasmic reticulum but does not perturb protein transport. Traffic 9(1), 94–102 (2008)

[6] Hu, J., Shibata, Y., Zhu, P.P., Voss, C., Rismanchi, N., Prinz, W.A., Rapoport, T.A., Blackstone, C.: A class of dynamin-like gtpases involved in the generation of the tubular er network. Cell 138(3), 549–61 (2009) 10.1016/j.cell.2009.05.025

[7] Orso, G., Pendin, D., Liu, S., Tosetto, J., Moss, T.J., Faust, J.E., Micaroni, M., Egorova, A., Martinuzzi, A., McNew, J.A., Daga, A.: Homotypic fusion of er membranes requires the dynamin-like gtpase atlastin. Nature 460(7258), 978–83 (2009) 10.1038/nature08280

[8] Chen, S., Novick, P., Ferro-Novick, S.: Er network formation requires a balance of the dynamin-like gtpase sey1p and the lunapark family member lnp1p. Nat Cell Biol 14(7), 707–16 (2012) 10.1038/ncb2523

[9] Wang, S., Tukachinsky, H., Romano, F.B., Rapoport, T.A.: Cooperation of the er-shaping proteins atlastin, lunapark, and reticulons to generate a tubular membrane network. Elife 5 (2016) 10.7554/eLife.18605

[10] Tikhomirova, M.S., Kadosh, A., Saukko-Paavola, A.J., Shemesh, T., Klemm, R.W.: A role for endoplasmic reticulum dynamics in the cellular distribution of microtubules. Proceedings of the National Academy of Sciences 119(15), 2104309119 (2022)

[11] Chen, S., Desai, T., McNew, J.A., Gerard, P., Novick, P.J., Ferro-Novick, S.: Lunapark stabilizes nascent three-way junctions in the endoplasmic reticulum. Proc Natl Acad Sci U S A 112(2), 418–23 (2015) 10.1073/pnas.1423026112

[12] Terasaki, M., Chen, L.B., Fujiwara, K.: Microtubules and the endoplasmic reticulum are highly interdependent structures. The Journal of cell biology 103(4), 1557–1568 (1986)

[13] Gao, G., Zhu, C., Liu, E., Nabi, I.R.: Reticulon and climp-63 control nanodomain organization of peripheral er tubules. PLOS Biology 17(8), 3000355 (2019)

[14] Schroeder, L.K., Barentine, A.E.S., Merta, H., Schweighofer, S., Zhang, Y., Baddeley, D., Bewersdorf, J., Bahmanyar, S.: Dynamic nanoscale morphology of the er surveyed by sted microscopy. J Cell Biol 218(1), 83–96 (2019) 10.1083/jcb.201809107

[15] Nixon-Abell, J., Obara, C.J., Weigel, A.V., Li, D., Legant, W.R., Xu, C.S., Pasolli, H.A., Harvey, K., Hess, H.F., Betzig, E., Blackstone, C., Lippincott-Schwartz, J.: Increased spatiotemporal resolution reveals highly dynamic dense tubular matrices in the peripheral er. Science 354(6311), 3928 (2016) 10.1126/science.aaf3928

[16] Holcman, D., Parutto, P., Chambers, J.E., Fantham, M., Young, L.J., Marciniak, S.J., Kaminski, C.F., Ron, D., Avezov, E.: Single particle trajectories reveal active endoplasmic reticulum luminal flow. Nature cell biology 20(10), 1118–1125 (2018)

[17] Porter, K.R., Yamada, E.: Studies on the endoplasmic reticulum. v. its form and differentiation in pigment epithelial cells of the frog retina. J Biophys Biochem Cytol 8(1), 181–205 (1960) 10.1083/jcb.8.1.181

[18] Tormey, J.M.: Fine structure of the ciliary epithelium of the rabbit, with particular reference to “infoldedmembranes,” “vesicles,” and the effects of diamox. J Cell Biol 17(3), 641–59 (1963) 10.1083/jcb.17.3.641

[19] Petrie, B.L., Graham, D.Y., Hanssen, H., Estes, M.K.: Localization of rotavirus antigens in infected cells by ultrastructural immunocytochemistry. J Gen Virol 63(2), 457–67 (1982) 10.1099/0022-1317-63-2-457

[20] Long, R.K.M., Moriarty, K.P., Cardoen, B., Gao, G., Vogl, A.W., Jean, F., Hamarneh, G., Nabi, I.R.: Super resolution microscopy and deep learning identify zika virus reorganization of the endoplasmic reticulum. Scientific Reports 10(1), 20937 (2020) 10.1038/s41598-020-77170-3

[21] Sparkes, I., Runions, J., Hawes, C., Griffing, L.: Movement and remodeling of the endoplasmic reticulum in nondividing cells of tobacco leaves. The Plant Cell 21(12), 3937–3949 (2009)

[22] Lin, C., Zhang, Y., Sparkes, I., Ashwin, P.: Structure and dynamics of er: minimal networks and biophysical constraints. Biophysical Journal 107(3), 763–772 (2014)

[23] Lin, C., Lemarchand, L., Euler, R., Sparkes, I.: Modeling the geometry and dynamics of the endoplasmic reticulum network. IEEE/ACM Transactions on Computational Biology and Bioinformatics 15(2), 377–386 (2015)

[24] Lin, C., White, R.R., Sparkes, I., Ashwin, P.: Modeling endoplasmic reticulum network maintenance in a plant cell. Biophysical Journal 113(1), 214–222 (2017)

[25] Euler, R., Lemarchand, L., Lin, C., Sparkes, I.: Modeling the geometry of the endoplasmic reticulum network. In: 20th Conference of the International Federation of Operational Research Societies IFORS 2014 (2014)

[26] Brown, A.I., Westrate, L.M., Koslover, E.F.: Impact of global structure on diffusive exploration of organelle networks. Scientific reports 10(1), 4984 (2020)

[27] Perkins, H.T., Allan, V.J., Waigh, T.A.: Network organisation and the dynamics of tubules in the endoplasmic reticulum. Scientific Reports 11(1), 16230 (2021)

[28] Pain, C., Kriechbaumer, V., Kittelmann, M., Hawes, C., Fricker, M.: Quantitative analysis of plant er architecture and dynamics. Nature communications 10(1), 984 (2019)

[29] Lu, M., Tartwijk, F.W., Lin, J.Q., Nijenhuis, W., Parutto, P., Fantham, M., Christensen, C.N., Avezov, E., Holt, C.E., Tunnacliffe, A., et al.: The structure and global distribution of the endoplasmic reticulum network are actively regulated by lysosomes. Science advances 6(51), 7209 (2020)

[30] Lu, M., Christensen, C.N., Weber, J.M., Konno, T., Läubli, N.F., Scherer, K.M., Avezov, E., Lio, P., Lapkin, A.A., Kaminski Schierle, G.S., et al.: Ernet: a tool for the semantic segmentation and quantitative analysis of endoplasmic reticulum topology. Nature Methods 20(4), 569–579 (2023)

[31] Garcia-Pardo, M.E., Simpson, J.C., O’Sullivan, N.C.: A novel automated image analysis pipeline for quantifying morphological changes to the endoplasmic reticulum in cultured human cells. BMC bioinformatics 22, 1–17 (2021)

[32] Pain, C., Kriechbaumer, V.: Defining the dance: quantification and classification of endoplasmic reticulum dynamics. Journal of Experimental Botany 71(6), 1757–1762 (2020)

[33] Bouchekhima, A.-N., Frigerio, L., Kirkilionis, .M.: Geometric quantification of the plant endoplasmic reticulum. Journal of microscopy 234(2), 158–172 (2009)

[34] Jerman, T., Pernuš, F., Likar, B., Špiclin, Ž.: Enhancement of vascular structures in 3d and 2d angiographic images. IEEE transactions on medical imaging 35(9), 2107–2118 (2016)

[35] Rumelhart, D.E., Hinton, G.E., Williams, R.J., et al.: Learning internal representations by error propagation. Institute for Cognitive Science, University of California, San Diego La … (1985)

[36] Ballard, D.H.: Modular learning in neural networks. In: Proceedings of the Sixth National Conference on Artificial Intelligence-volume 1, pp. 279–284 (1987)

[37] Sternberg: Biomedical image processing. Computer 16(1), 22–34 (1983) 10.1109/MC.1983.1654163

[38] Lee, T.-C., Kashyap, R.L., Chu, C.-N.: Building skeleton models via 3-d medial surface axis thinning algorithms. CVGIP: Graphical Models and Image Processing 56(6), 462–478 (1994)

[39] Shimodaira, H.: Improving predictive inference under covariate shift by weighting the log-likelihood function. Journal of statistical planning and inference 90(2), 227–244 (2000)

[40] Huang, J., Gretton, A., Borgwardt, K., Schölkopf, B., Smola, A.: Correcting sample selection bias by unlabeled data. Advances in neural information processing systems 19 (2006)

[41] Shorten, C., Khoshgoftaar, T.M.: A survey on image data augmentation for deep learning. Journal of big data 6(1), 1–48 (2019)

[42] Lenc, K., Vedaldi, A.: Understanding image representations by measuring their equivariance and equivalence. In: Proceedings of the IEEE Conference on Computer Vision and Pattern Recognition, pp. 991–999 (2015)

[43] Bekkers, E.J., Lafarge, M.W., Veta, M., Eppenhof, K.A., Pluim, J.P., Duits, R.: Roto-translation covariant convolutional networks for medical image analysis. In: Medical Image Computing and Computer Assisted Intervention–MICCAI 2018: 21st International Conference, Granada, Spain, September 16-20, 2018, Proceedings, Part I, pp. 440–448 (2018). Springer

[44] Wang, Y., Yao, Q., Kwok, J.T., Ni, L.M.: Generalizing from a few examples: A survey on few-shot learning. ACM computing surveys (csur) 53(3), 1–34 (2020)

[45] Rizve, M.N., Khan, S., Khan, F.S., Shah, M.: Exploring complementary strengths of invariant and equivariant representations for few-shot learning. In: Proceedings of the IEEE/CVF Conference on Computer Vision and Pattern Recognition, pp. 10836–10846 (2021)

[46] Jing, L., Tian, Y.: Self-supervised visual feature learning with deep neural networks: A survey. IEEE transactions on pattern analysis and machine intelligence 43(11), 4037–4058 (2020)

[47] Goyal, U., Blackstone, C.: Untangling the web: mechanisms underlying er network formation. Biochimica et Biophysica Acta (BBA)-Molecular Cell Research 1833(11), 2492–2498 (2013)

[48] Friedman, J.R., Webster, B.M., Mastronarde, D.N., Verhey, K.J., Voeltz, G.K.: Er sliding dynamics and er–mitochondrial contacts occur on acetylated microtubules. Journal of Cell Biology 190(3), 363–375 (2010)

[49] Puhka, M., Joensuu, M., Vihinen, H., Belevich, I., Jokitalo, E.: Progressive sheet-to-tubule transformation is a general mechanism for endoplasmic reticulum partitioning in dividing mammalian cells. Molecular biology of the cell 23(13), 2424–2432 (2012)

[50] Pizer, S.M., Amburn, E.P., Austin, J.D., Cromartie, R., Geselowitz, A., Greer, T., Haar Romeny, B., Zimmerman, J.B., Zuiderveld, K.: Adaptive histogram equalization and its variations. Computer vision, graphics, and image processing 39(3), 355–368 (1987)

[51] Walt, S., Schönberger, J.L., Nunez-Iglesias, J., Boulogne, F., Warner, J.D., Yager, N., Gouillart, E., Yu, T.: scikit-image: image processing in python. PeerJ 2, 453 (2014)

[52] Cohen, T., Welling, M.: Group equivariant convolutional networks. In: International Conference on Machine Learning, pp. 2990–2999 (2016). PMLR

[53] Ling, S.Z., Sharp, N., Jacobson, A.: Vectoradam for rotation equivariant geometry optimization. Advances in Neural Information Processing Systems 35, 4111–4122 (2022)

[54] Kingma, D.P., Ba, J.: Adam: A method for stochastic optimization. arXiv preprint arXiv:1412.6980 (2014)

[55] Wang, A., Yan, X., Wei, Z.: Imagepy: an open-source, python-based and platform-independent software package for bioimage analysis. Bioinformatics 34(18), 3238–3240 (2018)

[56] Hagberg, A., Swart, P., S Chult, D.: Exploring network structure, dynamics, and function using networkx. Technical report, Los Alamos National Lab.(LANL), Los Alamos, NM (United States) (2008)

[57] Samudre, A.: nERdy_data_v1 (2024) 10.6084/m9.figshare.25241458.v2

